# Perturbation of genes linked to common schizophrenia risk variants identifies cilia programs

**DOI:** 10.64898/2026.06.09.731172

**Authors:** Jiseok Lee, Hyunggyu Min, Cristine Casingal, Austin T. Ledford, Hannah Lee, Won Ma, Yangzhenyu Gao, Hanqian Mao, Eric S. McCoy, Lei Xing, Cindy Fang, Sang Ho Kwon, Mark J. Zylka, Keri Martinowich, Kristen R. Maynard, Stephanie C. Hicks, E. S. Anton, Hyejung Won

**Affiliations:** Neuroscience Center, University of North Carolina at Chapel Hill, Chapel Hill, NC; Department of Biostatistics, University of North Carolina at Chapel Hill, Chapel Hill, NC; Department of Biostatistics, Johns Hopkins Bloomberg School of Public Health, Baltimore, MD; Lieber Institute for Brain Development, Johns Hopkins Medical Campus, Baltimore, MD; The Solomon H. Snyder Department of Neuroscience, Johns Hopkins School of Medicine, Baltimore, MD, USA; Department of Psychiatry and Behavioral Sciences, Johns Hopkins School of Medicine, Baltimore, MD, USA; Johns Hopkins Kavli Neuroscience Discovery Institute, Baltimore, MD, USA

## Abstract

Schizophrenia (SCZ) is a common psychiatric disorder characterized by psychosis, emotional withdrawal, and cognitive deficits. Most SCZ risk variants reside in non-coding regions of the genome and are thought to influence disease risk by modulating gene regulation. However, the target genes, biological pathways, and cell types through which these variants exert their effects remain poorly understood. To address this gap, we employed *in vivo* CRISPR droplet sequencing (CROP-seq) in the postnatal mouse neocortex. We perturbed 12 SCZ risk genes previously linked to functionally validated risk variants, followed by single-cell RNA sequencing. We identified 3,031 differentially expressed genes (DEGs) that recapitulate transcriptional alterations observed in postmortem SCZ brains. Integrative analysis using DEG clustering, factor analysis, and gene regulatory network inference uncovered convergent gene programs with distinct biological functions and cell type specificity. Notably, ciliary transcriptional programs consistently emerged across analytical frameworks. The primary cilium is a neurocircuit modulating signaling organelle in neurons and glia that remains understudied in SCZ. Perturbation of key contributors to the ciliary transcriptional programs led to significant alterations in ciliary structure, suggesting that SCZ genetic risk factors may influence how brain cells sense and transduce extracellular signals through synapse-independent mechanisms. Together, this study provides the first *in vivo* characterization of the functional consequence of common variant architecture in SCZ and implicates ciliary dysfunction as a convergent downstream mechanism.

## INTRODUCTION

Schizophrenia (SCZ) is a mental disorder characterized by delusions, hallucinations, disorganized speech or behavior, and reduced emotional expression or motivation.^1^ SCZ affects 1 in 300 people worldwide according to estimates by the World Health Organization. SCZ typically onsets after late adolescence, though prior studies suggest developmental origin of the disorder.^2,3^ SCZ has a strong genetic influence with an estimated heritability of 80%.^4,5^ Genome-wide association studies (GWAS)^6,7^ revealed that SCZ risk is distributed across ∼300 genomic loci, whereas an exome sequencing study^8^ identified 32 genes with rare coding mutations, indicating that the disease risk is primarily carried by common variants. The vast majority of SCZ-associated common variants are located in the non-coding regions with elusive functional consequences. They are thought to influence gene regulation in a cell type-specific manner,^9^ making it crucial to link these variants to their target genes to uncover the biological mechanisms underlying SCZ.

To better understand the functional consequences of non-coding SCZ risk variants, we previously employed massively-parallel reporter assay (MPRA) to identify variants with empirically validated allelic regulatory activity.^10^ We then mapped these MPRA-validated variants to putative target genes using long-range chromatin interactions.^10^ A total of 272 genes interacting with 209 variants were identified using the chromatin loops. Notably, the putative target genes were enriched for transcriptional regulators, suggesting that their dysregulation could alter broader transcriptional networks. However, their functional roles in shaping transcriptional networks remain to be elucidated.

To understand how genes impacted by common variants contribute to the molecular pathology of SCZ, we applied CRISPR perturbation of 12 genes followed by single-cell transcriptomics in the developing mouse neocortex. While similar *in vivo* strategies have been used to study transcriptional programs affected by neurodevelopmental disorders^11–14^ such as autism spectrum disorder (ASD)^11^ and 22q11.2 deletion syndrome^14^, this is the first study linking common variant risk to SCZ mechanisms. Across perturbations, we identified ∼3,000 differentially expressed genes (DEGs), which closely recapitulated transcriptional changes observed in the brains of SCZ patients. These DEGs were distilled into convergent gene programs with distinct functional roles and cell type enrichment. While many of these programs converged on synaptic pathways previously implicated in SCZ, the ciliary program showed one of the strongest enrichment, despite being relatively understudied in SCZ pathophysiology. By perturbing key contributors to the ciliary transcriptional program, *Matr3* and *Scyl1*, we observed a significant alteration in primary cilia structure in neocortical cells. The changes in primary cilia structure may alter the way cortical cells access, correspond, and modulate the surrounding circuitry, thus leading to circuit malfunctions.

## RESULTS

### CRISPR perturbation of SCZ risk variant-associated genes in the mouse neocortex

To systematically dissect the transcriptional consequences of SCZ risk variants at single-cell resolution, we employed CRISPR droplet sequencing (CROP-seq),^15^ a high-throughput platform that enables CRISPR-based gene perturbation coupled with single-cell RNA sequencing (scRNA-seq) readout (**Figure 1A-B**). Among the 272 target genes mapped to MPRA-validated SCZ risk variants,^10^ we selected 12 genes encoding transcriptional regulators (e.g., transcription factors, DNA/RNA binding proteins, chromatin remodelers, nuclear proteins) to maximize our ability to capture transcriptional effects (**Table S1A**). We designed two guide-RNAs (gRNAs) per gene, targeting the beginning of the coding sequence for Cas9-mediated perturbation. These gRNAs were cloned into a modified CROP-seq vector expressing red fluorescent protein (RFP)^16^ for fluorescence-activated cell sorting (FACS), which is subsequently packaged into lentivirus. Since early developmental perturbations may reflect causal disease mechanisms rather than late-stage consequences, we injected the pooled lentiviral gRNA library into the lateral ventricles of postnatal day 0 (P0) Cas9 transgenic mice.^17^ This resulted in expression in the cerebral neocortex (**Figure S1A**), a key region implicated in SCZ.^18–20^ At P14, the neocortex was dissociated, RFP+ cells were sorted by FACS (**Figure S1B**), and processed for scRNA-seq. After quality control (**Methods**), a total of 95,495 cells representing the major cell types of the mouse neocortex were obtained (**Figure 1C**). Astrocytes and neurons were most abundant, followed by microglia, oligodendrocytes, and vascular cells (**Figure 1C**). Each major cell type was composed of various subtypes from the Allen Brain reference dataset^21^ (**Figure S1C**). We detected an average of ∼9,800 unique molecular identifier (UMI) counts and ∼2,800 unique genes per cell (**Figure S1D-E**). For gRNAs, we observed an average of ∼19 UMI counts and ∼10 unique gRNAs per cell, indicating a high multiplicity of infection (MOI; **Figure S1F-G**). On average, we collected ∼10,900 cells expressing each gRNA (**Figure S1H**).

**Figure 1.**
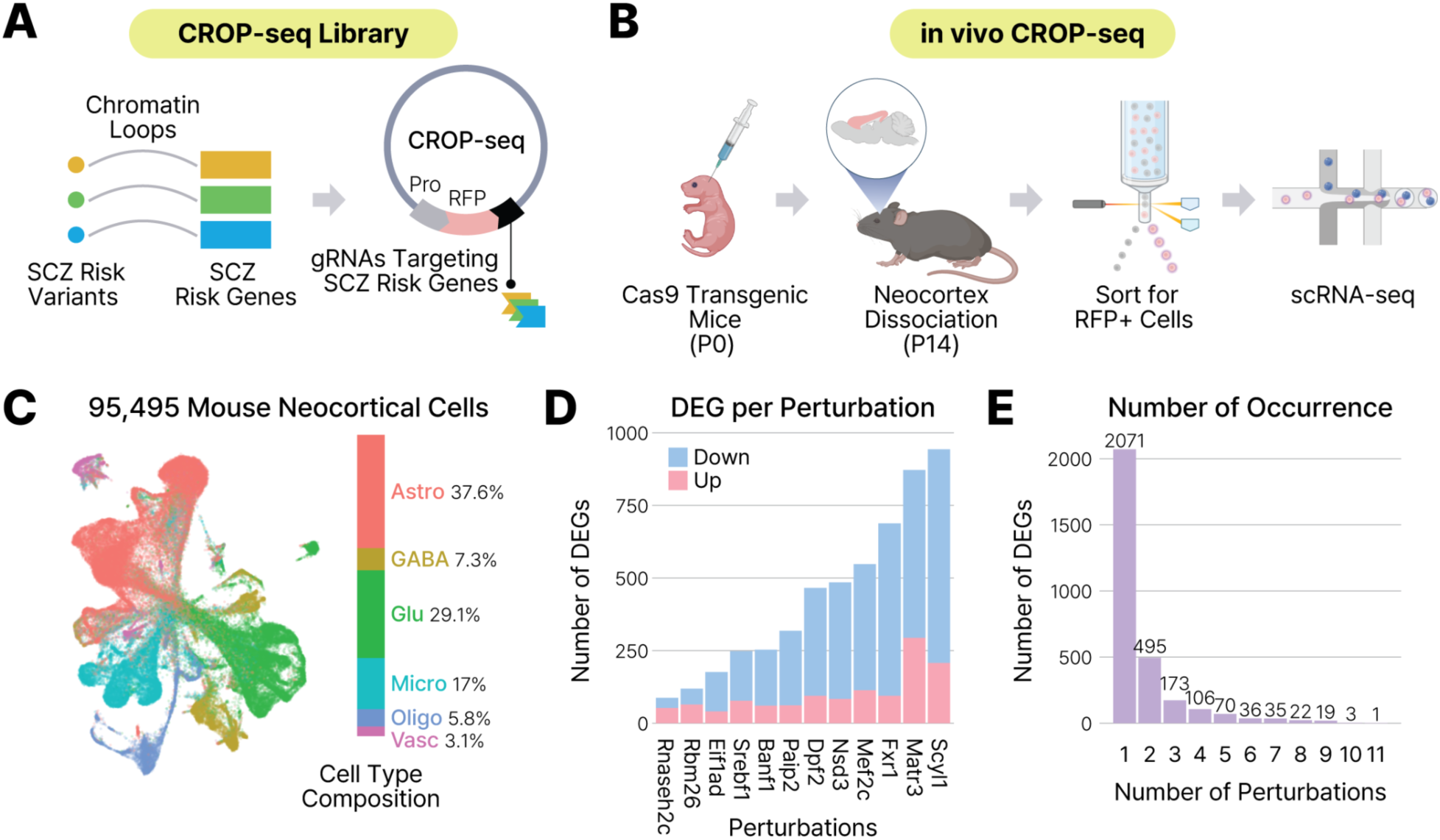
*In vivo* perturbation of SCZ risk genes in the mouse neocortex. **A.** SCZ risk genes were nominated based on chromatin interactions with empirically validated risk variants and perturbed using CROP-seq vectors encoding gene-targeting gRNAs. **B.** CROP-seq libraries were injected into Cas9 transgenic mice at P0. At P14, the neocortex was dissociated, and RFP+ cells expressing the library were isolated for scRNA-seq. **C.** Distribution of major cell types captured in the CROP-seq dataset. Astro, astrocytes; GABA, GABAergic neurons; Glu, glutamatergic neurons; Micro, microglia; Oligo, oligodendrocytes; Vasc, vascular cells. **D.** Number of DEGs for each perturbation, with colors indicating the proportions of downregulated and upregulated genes. **E.** Distribution of DEGs shared across perturbations.

To investigate transcriptional consequences of CRISPR perturbation, we carried out differential gene expression analysis using a negative binomial generalized linear model (GLM), followed by permutation analysis (**Methods, Figure S2, Table S1B**). We identified a total of 3,031 unique DEGs across the 12 perturbations (**Figure 1D, Table S1C**). Of the 3,031 DEGs, ∼65% were downregulated and ∼35% were upregulated (**Figure 1D**). Notably, ∼32% of DEGs (960 out of 3,031) were present in two or more perturbations (**Figure 1E**), suggesting molecular convergence of downstream transcriptional effects. We note, however, this could also be partly due to high-MOI gRNA expression (**Figure S1G**).

### Transcriptional signatures of SCZ risk gene perturbation

We first grouped DEGs as downregulated or upregulated, and characterized their cell type specificity and biological function (**Figure 2A**). All but three DEGs exhibited consistent directionality across perturbations. Downregulated and upregulated DEGs exhibited distinct cell type enrichment patterns, with downregulated DEGs showing strong neuronal enrichment and upregulated DEGs showing astrocytic enrichment (**Figure 2B**). Because ∼65% of DEGs were downregulated (**Figure 1D**), the overall DEG profile was dominated by neuronal enrichment (**Figure 2B**). This is consistent with transcriptional signatures observed in postmortem SCZ brains, which show downregulation of neuronal genes and upregulation of astrocytic genes.^22,23^

**Figure 2.**
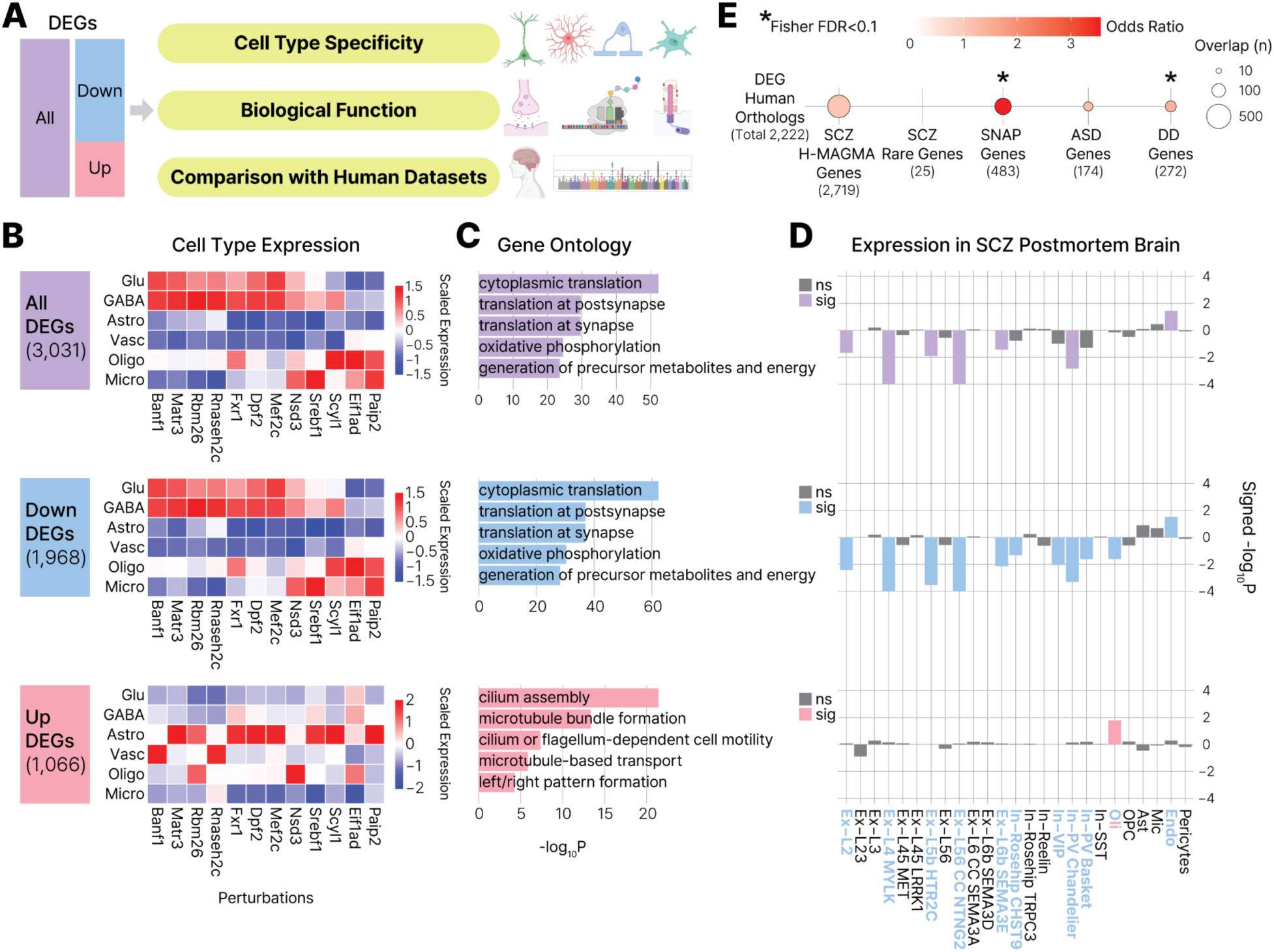
SCZ risk gene perturbation reveals disease-relevant transcriptional signatures. **A.** Schematic overview of the analytical framework. **B.** Scaled expression of all DEGs (top), downregulated DEGs (middle), and upregulated DEGs (bottom) across cell types for each perturbation. Numbers in parentheses indicate the total number of DEGs. **C.** Top 5 GO terms enriched in all DEGs (top), downregulated DEGs (middle), and upregulated DEGs (bottom). **D.** Expression of all DEGs (top), downregulated DEGs (middle), and upregulated DEGs (bottom) across 25 cell types in the postmortem prefrontal cortex (PFC) of individuals with SCZ. Colored bars and labels denote cell types in which DEGs are significantly dysregulated in SCZ. Negative and positive values indicate downregulation and upregulation, respectively. Ex-L2, layer 2 excitatory neuron; Ex-L23, layer 2/3 excitatory neuron; Ex-L3, layer 3 excitatory neuron; Ex-L4 MYLK, layer 4 MYLK+ excitatory neuron; Ex-L45 MET, layer 4/5 MET+ excitatory neuron; Ex-L45 LRRK1, layer 4/5 LRRK1+ excitatory neuron; Ex-L5b HTR2C, layer 5b HTR2C+ excitatory neuron; Ex-L56, layer 5/6 excitatory neuron; Ex-L56 CC NTNG2, layer 5/6 corticocortical NTNG2+ excitatory neuron; Ex-L6 CC SEMA3A, layer 6 corticocortical SEMA3A+ excitatory neuron; Ex-L6b SEMA3D, layer 6b SEMA3D+ excitatory neuron; Ex-L6b SEMA3E, layer 6b SEMA3E+ excitatory neuron; In-Rosehip CHST9, CHST9+ rosehip interneuron; In-Rosehip TRPC3, TRPC3+ rosehip interneuron; In-Reelin, reelin-expressing interneuron; In-VIP, vasoactive intestinal peptide-expressing interneuron; In-PV Chandelier, parvalbumin-expressing chandelier interneuron; In-PV Basket, parvalbumin-expressing basket interneuron; In-SST, somatostatin-expressing interneuron; Oli, oligodendrocyte; OPC, oligodendrocyte precursor cell; Ast, astrocyte; Mic, microglia; Endo, endothelial cell; Pericytes, pericyte. ns, non-significant; sig; significant. **E.** Enrichment of DEGs in human risk gene sets. Asterisks (*) indicate significant enrichment (FDR<0.1) by Fisher’s exact test. Numbers in parentheses indicate the number of genes used for each test.

Given this pronounced cell type difference, we conducted gene ontology (GO) analysis separately for downregulated and upregulated DEGs. Downregulated DEGs were enriched for GO terms related to translation, synapse, and mitochondrial function (**Figure 2C, middle**). The brain’s exceptionally high energetic demand to sustain neuronal activity,^24^ together with emerging evidence linking mitochondrial dysfunction to SCZ,^23,25–27^ underscores the functional relevance of these pathways. In contrast, upregulated DEGs showed selective enrichment for ciliary function (**Figure 2C, bottom**).

To assess the relevance of these transcriptional signatures in human disease, we compared CROP-seq DEGs with postmortem single-nucleus RNA sequencing (snRNA-seq) dataset from individuals with SCZ.^28^ Mouse CROP-seq data recapitulated cell type-specific transcriptional alterations observed in human postmortem SCZ samples, with downregulated DEGs broadly decreased across excitatory and inhibitory neuronal subtypes and upregulated DEGs selectively increased in oligodendrocytes (**Figure 2D**).

Finally, we tested whether downstream transcriptional programs of SCZ common variants are enriched for genetic risk for neurodevelopmental and psychiatric disorders by assessing overlap between CROP-seq DEGs with human disease-relevant gene sets (**Figure 2E**). We observed significant enrichment in two datasets. First, we observed enrichment for co-regulated synaptic neuron and astrocytic program (SNAP) genes implicated in SCZ,^29^ indicating that coordinated neuron-glia transcriptional programs previously linked to SCZ pathophysiology can be driven by common variants. Second, CROP-seq DEGs were enriched for genes harboring rare variants associated with developmental disorders (DD),^30^ suggesting convergence between neurodevelopmental disorders and SCZ despite their distinct genetic architectures: DD driven by rare, high-impact protein-truncating mutations and SCZ by common variants with small individual effect sizes.^3^

### Common SCZ risk genes converge on core biological programs

To gain deeper biological insight into the transcriptomic changes induced by SCZ risk gene perturbations, we employed two independent approaches to derive gene programs with distinct biological functions: (1) hierarchical clustering of DEGs and (2) consensus non-negative matrix factorization (cNMF).

In the first approach, we performed hierarchical clustering of DEGs based on the similarity of GLM coefficients, which indicate direction and magnitude of change across perturbations (**Figure 3A**), a strategy commonly used to derive gene programs.^31,32^ We selected the number of clusters based on the silhouette score (**Figure S3A**) and gene ontology analysis, balancing functional diversity with minimal redundancy. This analysis identified five gene programs with distinct functional annotations, encompassing neuroarchitecture (Program 1), synapse (Program 2), cilia (Program 3), translation (Program 4), and catabolism (Program 5) (**Figure 3B, Table S2A**). Consistent with the aggregated DEG analysis (**Figure 2C**), mitochondria-related GO terms were enriched in the neuroarchitecture and synapse programs. Notably, similar programs have been reported following perturbation of ASD risk genes in human brain organoids,^33^ suggesting convergence of downstream biological pathways across psychiatric and neurodevelopmental disorders.

**Figure 3.**
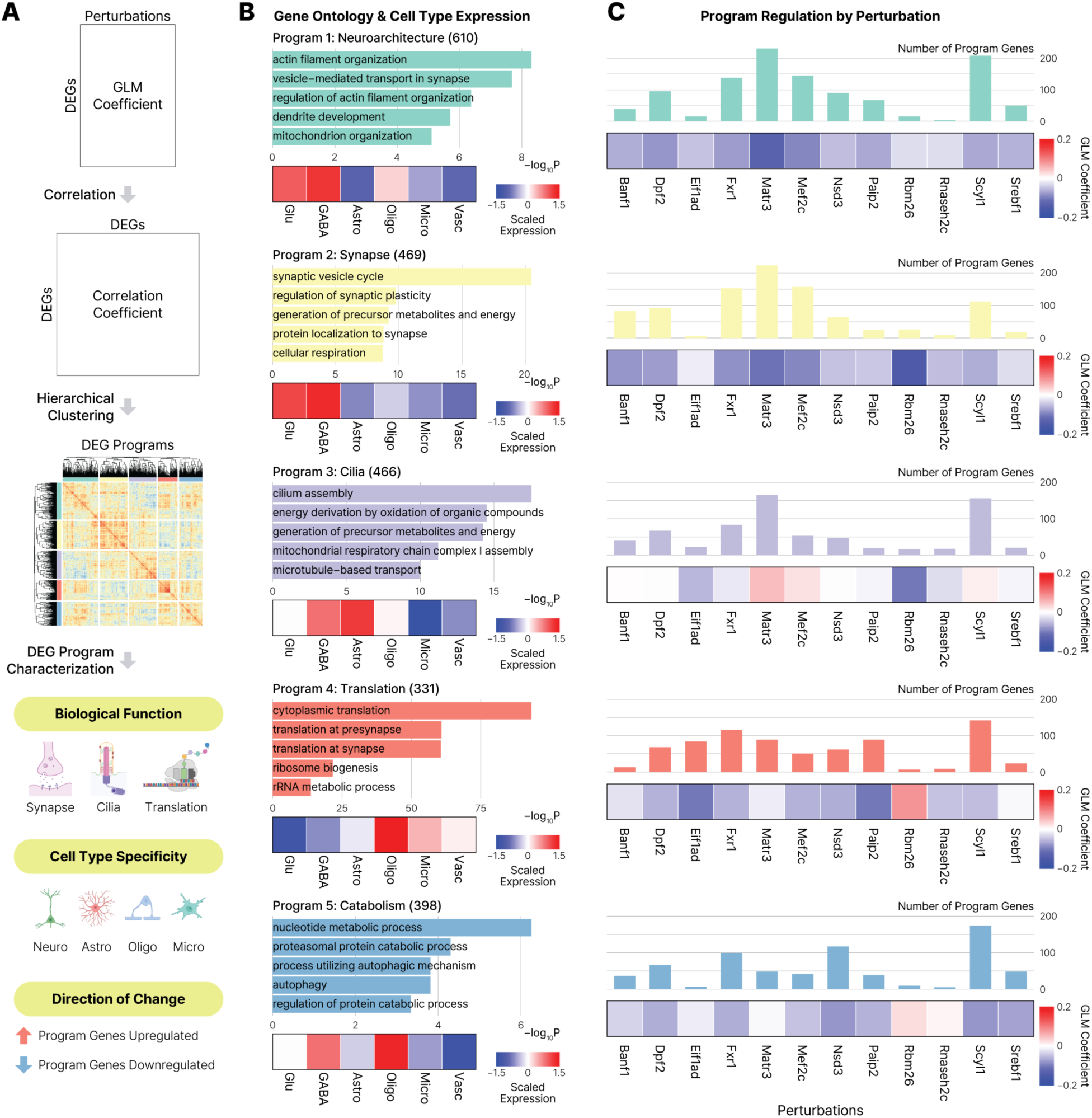
Identification of core biological programs downstream of SCZ risk genes. **A.** Schematic overview of DEG-based gene program identification. DEGs were grouped based on similarity of GLM coefficients, followed by hierarchical clustering to define gene programs. **B.** Functional and cell type characterization of DEG programs. For each program, the top 5 GO terms (bar graphs) and scaled expression across major cell types (heatmaps) are depicted. Numbers in parentheses indicate the number of genes in each program. **C.** Regulation of gene programs across perturbations. Bar graphs indicate the number of DEGs per program affected by each perturbation. Heatmaps show the corresponding GLM coefficients, reflecting the direction of change for perturbation-specific DEGs.

We next examined cell type specificity of these programs (**Figure 3B**; heatmaps, **Figure S3B**). Programs 1 (neuroarchitecture) and 2 (synapse) were predominantly enriched in neurons, while the remaining programs showed greater enrichment in glia: Program 3 (cilia) in astrocytes, and Programs 4 (translation) and 5 (catabolism) in oligodendrocytes. However, subsets of genes within each program were expressed across multiple cell types, indicating that these programs are enriched in, but not restricted to, specific cell types (**Figure S3B**). These results suggest that, despite the predominant enrichment of SCZ risk variants in neuronal enhancers,^3,6,9^ the downstream transcriptional consequences extend beyond neurons.^29^

Finally, we assessed how each program was regulated across perturbations (**Figure 3C**; heatmaps, **Figure S3C**). Most programs were predominantly downregulated across the 12 perturbations, while Program 3 (cilia) was frequently upregulated (**Figure 3C**, **Figure S3C**). Each program was shaped by multiple perturbations rather than a few drivers (**Figure 3C**; bar plots), suggesting convergence of SCZ risk genes onto shared functional pathways.

### cNMF-based transcriptional programs recapitulate SCZ-associated cellular and molecular signatures

To identify broader transcriptional programs regulated by perturbations beyond those captured by DEG-based analysis, we applied cNMF^34^ (**Figure 4A, Figure S4A, Methods**). We first decomposed the gene expression matrix into *k*=300 factors and subsequently refined this set to 118 perturbation-associated factors by selecting those enriched for DEGs. These factors were then clustered into 8 biologically interpretable programs based on GO analysis (hereafter referred to as P0-P7; **Figure 4B, Table S2B**).

**Figure 4.**
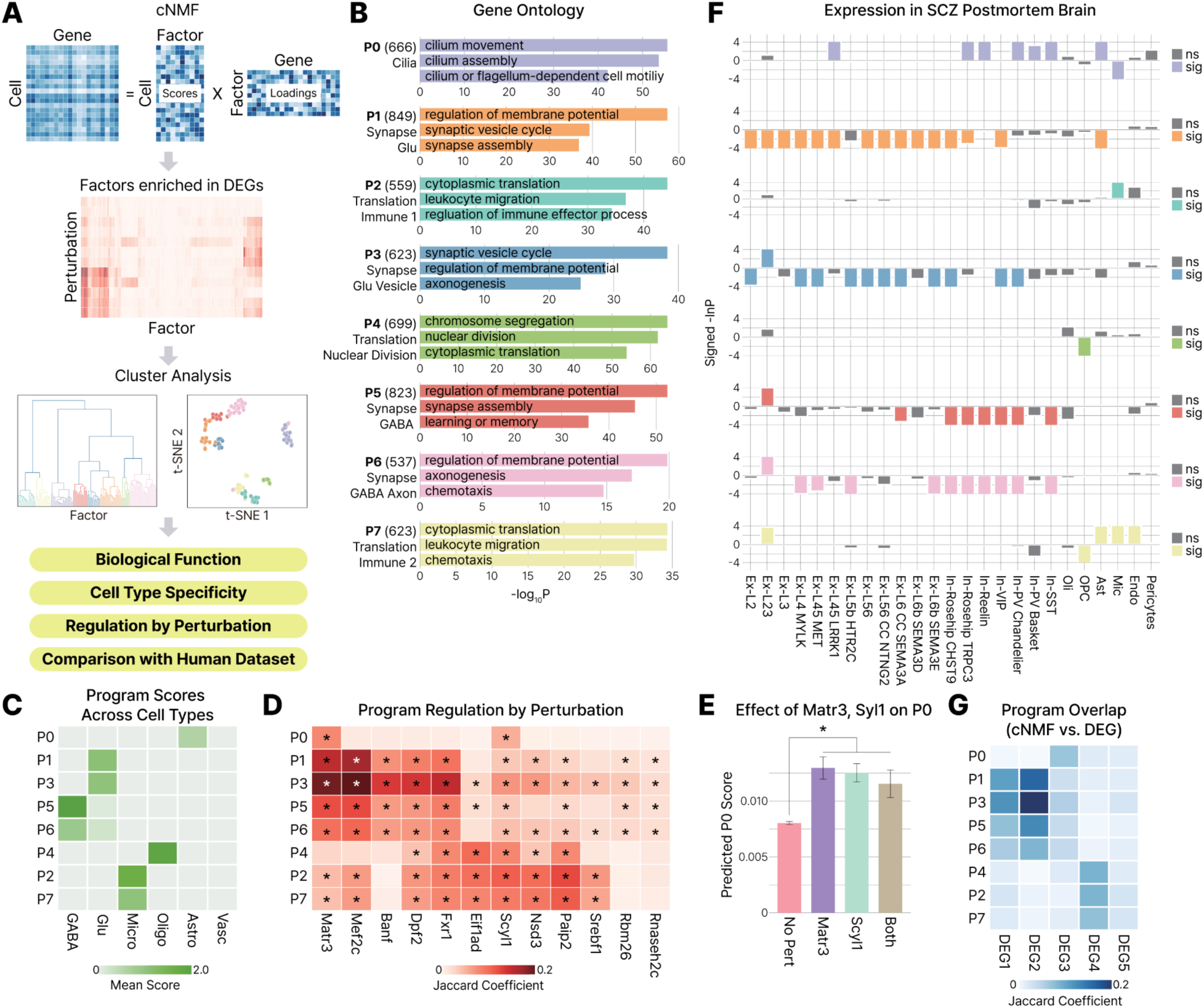
cNMF-derived programs capture cellular and molecular signatures downstream of SCZ risk genes. **A.** Schematic overview of cNMF program identification and downstream analysis. **B.** Eight cNMF programs and their top enriched GO terms. Numbers in parentheses indicate the number of genes assigned to each program. **C.** Program scores across cell types. For each program and cell type, cNMF scores were averaged across constituent factors and then across cells. **D.** Regulation of gene programs across perturbations. Asterisks (*) indicate significant overlap by Fisher’s exact test (FDR<0.05). **E.** Predicted cNMF P0 scores stratified by *Matr3* and *Scyl1* perturbation status. Asterisk (*) indicates significant difference relative to unperturbed cells (P<0.05). **F.** Program-level expression changes across 25 cell types in the postmortem SCZ PFC. Colored bars denote cell types in which program genes are significantly dysregulated in SCZ. **G.** Overlap between cNMF programs and DEG programs.

Consistent with DEG-based analysis, cNMF identified a cilia program (P0), multiple synapse-related programs (P1, P3, P5, and P6), and translation-related programs (P2, P4, and P7). Despite partial functional overlap, these programs showed distinct biological functions and cell type enrichment (**Figure 4C**). For example, P1 and P3 showed the highest cNMF scores in glutamatergic neurons, whereas P5 and P6 were most enriched in GABAergic neurons. P2 and P7 were associated with immune-related functions and enriched in microglia. P0 was enriched in astrocytes, consistent with the DEG-based cilia program (**Figure 3B**).

To determine how individual perturbations influenced each cNMF program, we tested the overlap between cNMF programs and perturbation-specific DEGs. All perturbations influenced at least four programs, and each program was affected by multiple perturbations (**Figure 4D**). In contrast, P0 was selectively affected by *Matr3* and *Scyl1* perturbations. Given the selective regulation of P0 by *Matr3* and *Scyl1*, we next sought to resolve the combinatorial effects of these perturbations. We modeled P0 scores as a function of perturbation status (**Figure 4E**). Perturbation of either *Matr3* or *Scyl1* significantly increased P0 scores compared to unperturbed cells, but we did not observe a synergistic effect. Together, these results suggest that *Matr3* and *Scyl1* are key drivers of the ciliary transcriptional program.

We next examined how cNMF programs were altered in postmortem SCZ brains, analogous to the DEG-based analysis (**Figure 4F**). cNMF programs exhibited cell type-specific dysregulation consistent with SCZ molecular pathology. Neuronal programs, including glutamatergic (P1, P3) and GABAergic (P5, P6) programs, were broadly downregulated in excitatory and inhibitory neuronal subtypes. In contrast, immune-related programs (P2, P7) were upregulated in microglia and astrocytes. The cilia program (P0) was upregulated in astrocytes and GABAergic neurons. The translation program (P4), which showed the strongest enrichment in oligodendrocytes, was specifically downregulated in oligodendrocyte precursor cells.

Finally, comparison of DEG-and cNMF-derived programs revealed substantial functional overlap between the two approaches (**Figure 4G**). Collectively, these results demonstrate that cNMF identifies additional fine-grained transcriptional programs relevant to SCZ molecular pathology.

### Extending SCZ-related gene regulatory network

Beyond defining gene programs perturbed by SCZ risk genes, we sought to uncover extended gene regulatory networks that could provide broader mechanistic insight. To this end, we applied pySCENIC^35^ to our CROP-seq data to infer ‘regulons’ (**Figure 5A**), or transcription factor (TF)-centered gene regulatory modules. Each regulon consists of a TF and its predicted target genes, inferred based on gene co-expression and refined by enrichment of the corresponding TF binding motif.

**Figure 5.**
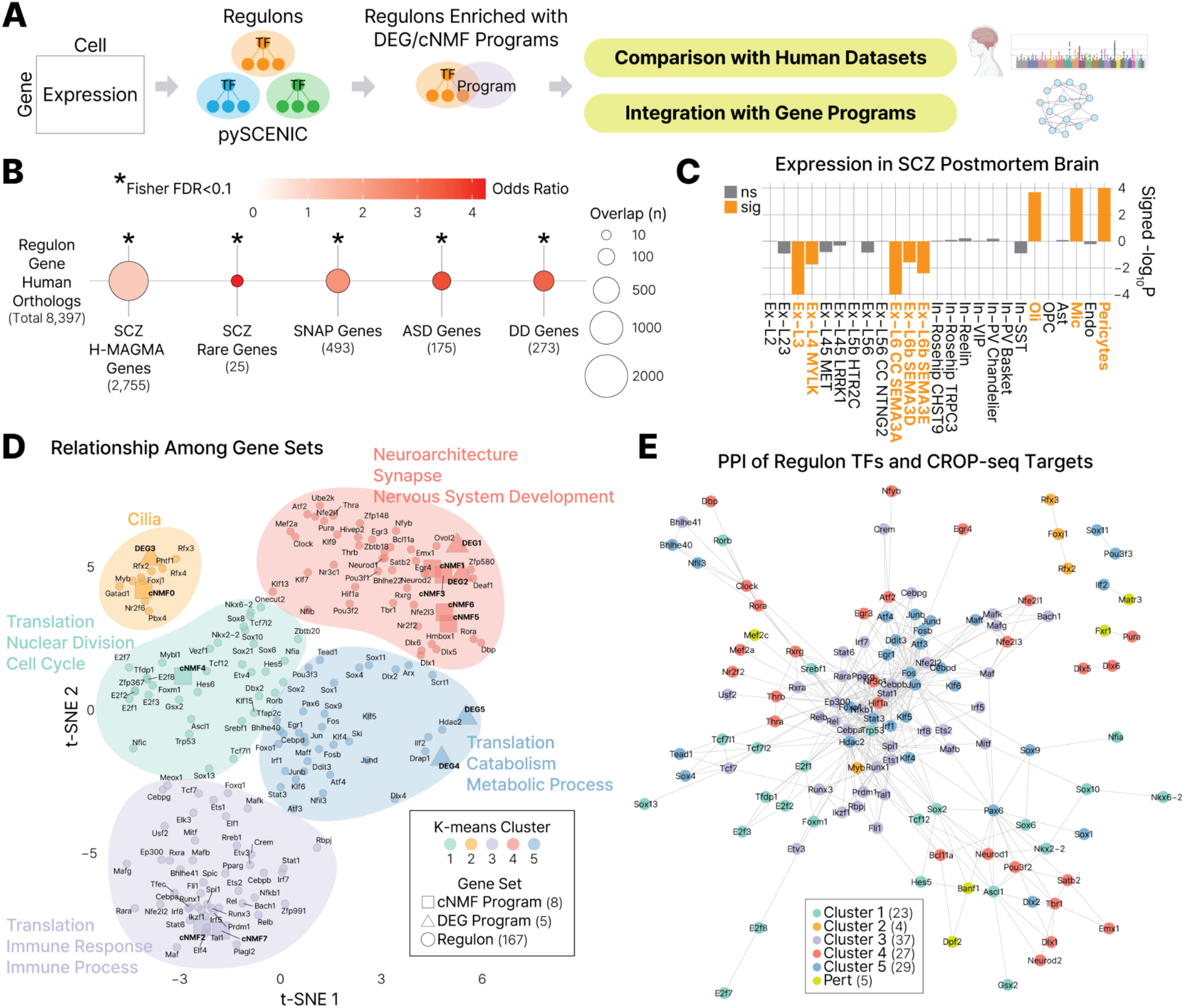
Regulon analysis reveals SCZ-associated gene regulatory networks. **A.** Schematic overview of regulon identification using psySCENIC, followed by enrichment testing and network integration with gene programs. **B.** Enrichment of regulon genes in human risk gene sets. Asterisks (*) indicate significant enrichment (FDR<0.1) by Fisher’s exact test. Numbers in parentheses indicate the number of genes used for each test. **C.** Expression of regulon genes across 25 cell types in the postmortem SCZ PFC. Colored bars and labels denote cell types in which regulon genes are significantly dysregulated in SCZ. **D.** Relationships among gene sets, including 5 DEG programs, 8 cNMF programs, and 167 regulons, visualized by dimensionality reduction and clustering. Five major clusters with distinct biological functions are identified. **E.** PPI network of regulon TFs and CROP-seq perturbation targets.

We identified 167 regulons encompassing 9,209 genes that were significantly enriched in either the DEG or cNMF programs (**Table S2C**). These regulons were enriched for SCZ genetic risk factors, including genes implicated by both common^2,6^ and rare^8^ variants (**Figure 5B**), suggesting that genes embedded within regulatory networks perturbed by SCZ risk genes also contribute to the genetic architecture of the disorder. Regulons were also enriched for SNAP genes,^29^ suggesting that these regulatory modules capture transcriptional programs altered in postmortem SCZ brains (**Figure 5B**). Furthermore, regulons were enriched for genes harboring rare variants associated with ASD^36^ and DD^30^ (**Figure 5B**), indicating that genes with key neurodevelopmental functions are central to SCZ-associated regulatory networks. Consistent with these findings, regulons were broadly downregulated in excitatory neuronal subtypes and upregulated in glia in postmortem SCZ brains (**Figure 5C**), mirroring the expression patterns of DEGs (**Figure 2D**) and cNMF programs (**Figure 4F**).

To provide an integrated view of these gene sets (DEG programs, cNMF programs, and regulons), we visualized their relationships by applying dimensionality reduction to a distance matrix followed by clustering (**Figure 5D**). This analysis revealed five major clusters, each comprising at least one DEG or cNMF program together with associated regulons, collectively representing distinct biological functions. In addition to the 12 perturbed risk genes, regulon TFs emerging from this analysis may act as upstream regulators of the transcriptional programs implicated in SCZ.

We next examined whether these TFs participate in protein-protein interaction (PPI) networks. Indeed, 120 regulon TFs and 5 perturbed risk genes formed interconnected PPI networks (**Figure 5E**). Notably, Foxj1, Rfx2, and Rfx3, previously reported as key regulators of ciliogenesis,^37,38^ were associated with cilia programs (**Figure 5D**) and exhibited protein-level interactions.

### Linking SCZ common risk to ciliary dysfunction

Across analyses, perturbation of SCZ risk genes consistently highlighted cilia-related functions (**Figure 2C, 3B, 4B, 5D**). Cilia are centrosome-derived, microtubule-based organelles that protrude from the cell surface. All neurons and astrocytes possess a single non-motile cilium,^39^ termed the primary cilium, which functions as a sensory antenna to detect extracellular cues and mediate intracellular signaling. A wide range of neurotransmitter and neuromodulator receptors, as well as synaptic proteins, localize to the primary cilium.^40–42^ Emerging evidence links ciliary dysfunction to SCZ, including altered expression of ciliary genes in postmortem brains^43^ and aberrant cilia formation in patient-derived olfactory neuronal precursors.^44^

However, the role of ciliary function in SCZ remains incompletely understood. We therefore sought to further interrogate its role in SCZ molecular pathology. **Figure 6A** summarizes 55 perturbation-associated DEGs with strong ciliary association (**Table S3A**), curated based on subcellular localization from prior studies.^45,46^ In addition, 109 DEGs overlapped with mouse neuronal cilia proteome (**Table S3B**).^40^ Perturbation of *Matr3* and *Scyl1* contributed the largest number of DEGs within the cilia program (**Figure 3C**), and their associated DEGs exhibited significantly higher cilia association scores compared to background genes (**Figure 6B**). We therefore used *Matr3* and *Scyl1* DEGs to assess the relevance of ciliary signatures in SCZ.

**Figure 6.**
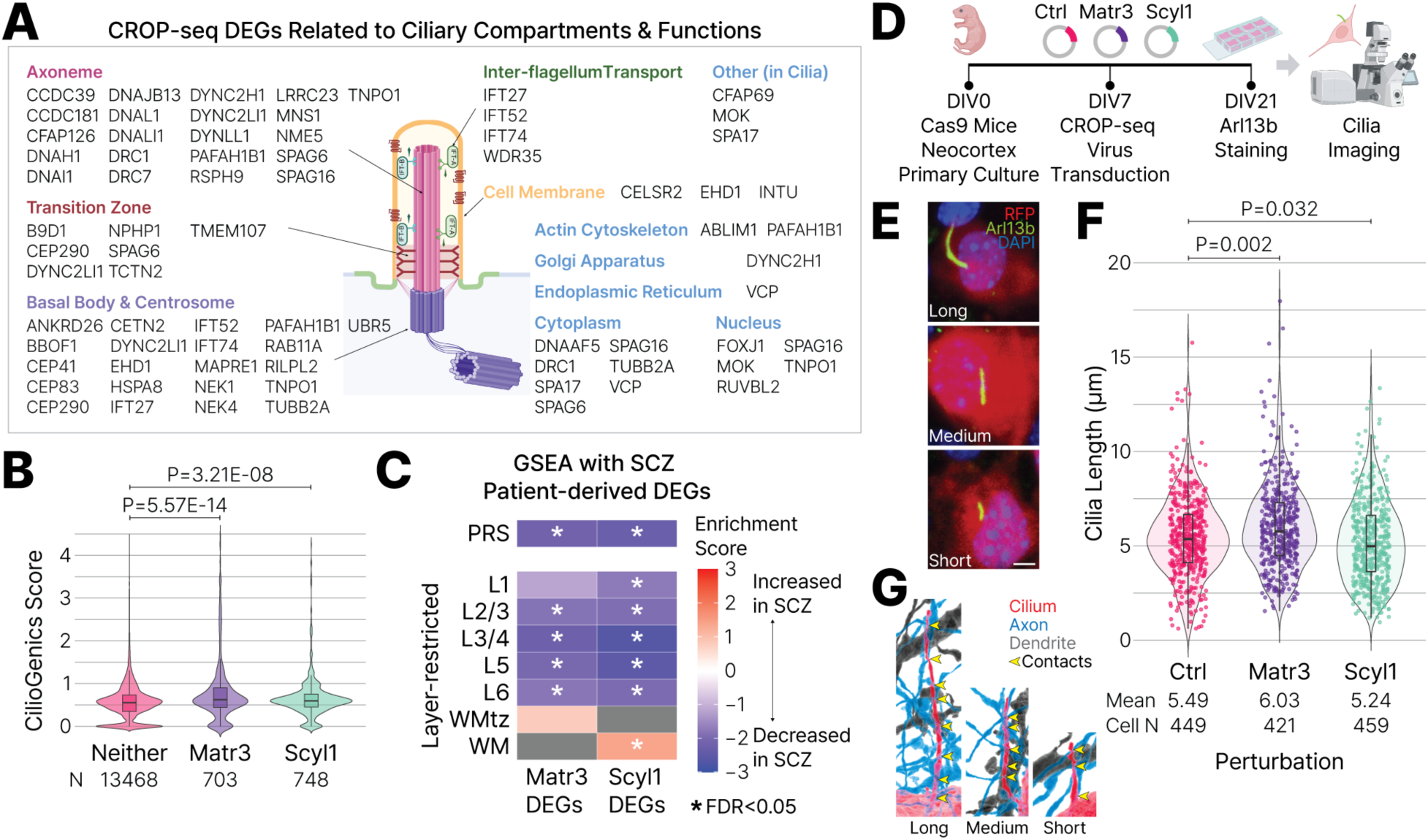
SCZ common risk is associated with ciliary dysfunction. **A.** DEGs with strong ciliary association and their subcellular localization. **B.** Distribution of CilioGenics scores (indicating cilia association) for background genes (Neither), *Matr3* DEGs, and *Scyl1* DEGs. P-values calculated by Wilcoxon rank-sum test. **C.** GSEA of *Matr3* and *Scyl1* DEGs using PRS-associated and layer-restricted gene expression profiles from postmortem SCZ brains. Positive and negative enrichment scores indicate increased and decreased expression with increasing PRS or SCZ condition, respectively. Asterisks (*) indicate significant enrichment (FDR<0.05). Grey boxes indicate cases in which GSEA failed to calculate P-values. L1, layer 1; L2/3, layers 2-3; L3/4, layers 3-4; L5, layer 5; L6, layer 6; WMtz, white matter transition zone; WM, white matter. **D.** Schematic overview of the cilia imaging experiments following *Matr3* and *Scyl1* perturbation. **E.** Representative examples of primary cilia classified as long, medium, and short. Scale bar, 5 μm. **F.** Distribution of cilia length in control, *Matr3*-, and *Scyl1*-perturbed cells. P-values calculated by Wilcoxon rank-sum test. **G.** Schematic illustrating the relationship between cilia length and contacts with adjacent axons and dendrites.

To determine whether these DEGs reflect common variant-mediated risk, we performed gene set enrichment analysis (GSEA) using polygenic risk score (PRS)-ranked transcriptomic data,^23^ testing for enrichment among genes up-or down-regulated with increasing polygenic burden. We found that both *Matr3* and *Scyl1* DEGs were significantly enriched among genes downregulated with increasing polygenic risk (**Figure 6C, top**).

We next leveraged spatial transcriptomic datasets^23^ to determine whether cilia-related programs are dysregulated in specific spatial domains. Using cortical layer-resolved transcriptomic data,^23^ in which genes were ranked by the direction and significance of their expression change in SCZ, we found that *Matr3* and *Scyl1* DEGs were significantly enriched among genes downregulated across layers 2-6 (**Figure 6C, bottom**). In addition, *Scyl1* DEGs were downregulated in layer 1 but upregulated in white matter.

Collectively, our analyses suggest that perturbation of genes linked to common SCZ risk variants induces cilia-associated transcriptional changes. Notably, these programs were significantly downregulated with increasing PRS, indicating convergence of genetic risk on ciliary dysfunction. This dysregulation was observed across cortical layers, suggesting a widespread cortical effect rather than spatially restricted alterations.

### Perturbation of the cilia program alters primary cilia length in mouse neocortical cells

Our findings implicate ciliary dysfunction as a potential downstream mechanism linking genetic risk to SCZ pathology. However, the extent to which perturbation-induced transcriptional changes translate into altered ciliary phenotypes remains unclear. We therefore selected two SCZ risk genes, *Matr3* and *Scyl1*, for experimental validation based on their strong contributions to the ciliary transcriptional programs upon perturbation (**Figures 3C, 4D-E**).

*MATR3* encodes Matrin-3 (MATR3), a nuclear scaffolding protein involved in chromatin organization through interactions with scaffold/matrix attachment regions (S/MAR sequences), chromatin remodelers, and chromatin architectural proteins.^47^ MATR3 is also a DNA-and RNA-binding protein that regulates transcription, DNA damage response, RNA stability, and splicing.^47^ Furthermore, MATR3 can localize to the cytoplasm, where it associates with the translation machinery.^47^ Mutations in *MATR3* have been implicated in amyotrophic lateral sclerosis (ALS) and neurodevelopmental disorders.^47^ In addition, a SCZ risk locus has been identified as a putative repressor of *MATR3*.^48^

*SCYL1* encodes SCY1-like pseudokinase 1 (SCYL1), a catalytically inactive kinase involved in Golgi apparatus-mediated intracellular protein trafficking through interactions with the coat protein complex I (COPI) coatomer.^49,50^ This is notable given the close functional and spatial relationship between the Golgi apparatus and the primary cilium.^51^ In addition to its role in protein trafficking, SCYL1 has been identified as a transcriptional regulator of human telomerase reverse transcriptase (TERT) and DNA polymerase beta (POLB).^49,52^ Mutations in *SCYL1* are associated with peripheral neuropathy, cerebellar atrophy, ataxia,^53,54^ and *Scyl1*-deficient mice exhibit features of ALS.^49,55^

To assess the effects of perturbing *Matr3* and *Scyl1* on ciliary morphology, we generated lentiviral gRNA libraries targeting each gene, along with a negative control library.^56^ These libraries were transduced into primary neocortical cultures derived from P0 Cas9 transgenic mice^17^ at 7 days in vitro (DIV7) (**Figure 6D**). At DIV21, cultures were fixed and immunostained for Arl13b, a canonical ciliary marker, and imaged by confocal microscopy (**Figure 6D**). We quantified primary cilia length in RFP+ cells expressing the gRNA libraries (**Figure 6E**). *Matr3* perturbation resulted in a significant increase in primary cilia length compared to control, whereas *Scyl1* perturbation resulted in a significant decrease (**Figure 6F, Figure S5A**). In contrast, analysis of cilia shape (smooth, irregular, branched) revealed no significant differences across perturbation groups (**Figure S5B-C**). Primary cilia length directly correlates with the number of functional contacts with adjacent axons, dendrites, and glial processes, reflecting its signaling capacity (**Figure 6G**). Together, these findings suggest that perturbation of key contributors to ciliary transcriptional programs remodels ciliary structure and may alter how neurons and astrocytes receive, send, and process extracellular signals.

## DISCUSSION

Using *in vivo* CROP-seq in the mouse neocortex, we characterized transcriptional changes induced by perturbation of 12 SCZ risk genes linked to common variants and identified convergent gene programs encompassing neuronal and synaptic functions, as well as mitochondrial activity, metabolism, translation, cell cycle regulation, immune responses, and ciliary function. These findings align with a growing body of evidence implicating diverse biological processes in SCZ pathophysiology.^25–27^

Although we employed Cas9-mediated gene knockout—an approach that more closely models rare, high-impact mutations than the subtle effects of common variants—common variants are thought to influence disease risk via gene regulatory mechanisms. Perturbation of their target genes therefore provide a tractable strategy to interrogate transcriptional programs through which polygenic risk is mediated. Consistent with this framework, transcriptional signatures induced by CROP-seq recapitulate those observed in postmortem SCZ brains. Moreover, the high MOI context, in which cells undergo multiple perturbations simultaneously, may partially model the polygenic and combinatorial nature of common variants.

Among transcriptional programs identified, ciliary functions were consistently enriched, motivating focused investigation of the primary cilium—an antenna-like organelle present in nearly all brain cells.^39,57^ Perturbation of *Matr3* and *Scyl1*, the strongest contributors to ciliary programs, resulted in significant changes in primary cilia length in mouse neocortical cells, indicating that transcriptional dysregulation identified by CROP-seq corresponds to functional changes in ciliary structure.

The length and shape of primary cilia directly correlate with the extent of access they have to the surrounding brain circuitry.^39,58^ Cortical neuronal primary cilia can make gap-junction contacts, axo-ciliary synapses, or tetrapartite synapses with the neural circuitry.^59^ Changes in primary cilia length as a result of disrupted *Matr3* and *Scyl1* activity can change primary cilia connectome, thus changing the way primary cilia access the neural circuitry. Although MATR3, a nuclear matrix protein, and SCYL1, a regulator of intracellular trafficking, likely affect ciliogenesis and maintenance through distinct mechanisms, their perturbation converges on a shared ciliary endophenotype that may destabilize the modulatory influence exerted by primary cilia on surrounding neural circuits.

Primary cilia signaling is increasingly recognized as a vulnerable nodal point in neuropsychiatric disorders. Expression of cilia-associated genes is significantly dysregulated in the frontal cortex of individuals with major psychiatric disorders including SCZ, ASD, bipolar disorder, and depression.^43^ Copy number variation of the complement component 4A (C4A) gene confers significant SCZ risk and has been linked to disruption of primary cilia-related processes in excitatory neurons.^60^ Furthermore, high-confidence ASD genes converge in expression, localization, and function at cilia, and patients carrying pathogenic variants in these genes exhibit cilia-related co-occurring conditions and biomarkers of disrupted ciliary function.^42^ Likewise, disruption of ciliary function in excitatory and inhibitory neurons can lead to altered synaptic connectivity and excitatory-inhibitory balance.^61–63^ Recent studies also revealed that novel axo-ciliary synapses can regulate the epigenetic state of post-synaptic neurons.^64^ Collectively, these findings highlight primary cilia as a critical integrative hub in neural circuit regulation and disorders. In this study, an unbiased functional interrogation of SCZ risk genes converged on cilia pathways as a point of vulnerability. Perturbation of SCZ risk genes that contribute to ciliary programs altered cilia structure, providing a mechanistic link between common variant risk and neural circuit function through modulation of cilia-dependent signaling.

### Limitations of the study

This study has several limitations. First, functional analyses were restricted to 12 SCZ risk genes, and expansion to a broader set of risk genes will be necessary for a more comprehensive understanding of common variant-mediated effects. Second, analyses were conducted at a single developmental time point (P14) and focused on one brain region (neocortex). Future studies incorporating additional risk genes, developmental stages, and brain regions will provide a more comprehensive understanding of SCZ pathophysiology. Third, experimental validation of ciliary phenotypes was limited to morphological measurements, such as cilia length, and did not assess potential alterations in ciliary signaling cascades such as Ca^2+^ or ciliary synaptic contacts. Incorporating functional readouts of ciliary signaling and ciliary synapses will be important to more fully define the role of primary cilia in circuit malfunction associated with SCZ pathophysiology. Finally, transcriptomic analyses were performed across all cells rather than individual cell types due to limited statistical power upon stratification. Nonetheless, gene programs derived from pooled analyses exhibited robust cell type specificity in both mouse and postmortem SCZ brains, supporting the utility of this approach.

## RESOURCE AVAILABILITY

### Lead contact

Any inquiries about the project, analytical results, or other information should be directed to Hyejung Won.

### Data and code availability

Raw and Cell Ranger-processed scRNA-seq data from CROP-seq are available under GEO accession number GSE334303. Custom codes used to analyze CROP-seq data are available on our GitHub page (https://github.com/thewonlab/SCZ_CROPseq).

## ACKNOWLEDGEMENTS

We thank the members of the Won lab for helpful discussions and comments on this manuscript. We thank Dr. Benjamin D. Philpot and Dr. Matthew C. Judson (UNC Neuroscience Center) for assistance with setting up the neonatal injections. We thank Dr. Xin Jin (Scripps Research) for sharing the single-cell dissociation protocol. We thank Dr. Lucas Ferreira (Harvard Medical School) for assistance with the Cell Ranger pipeline. We thank Dr. Timothy Barry (University of Pennsylvania) for discussions regarding differential expression analysis. We thank Dr. Jing Zhang and Siwei Xu for assistance with the SCENIC pipeline. We also acknowledge technical support from the UNC Lenti-shRNA Core, UNC Division of Comparative Medicine, UNC Advanced Analytics Core (Center for Gastrointestinal Biology and Disease grant P30DK034987), and the UNC High Throughput Sequencing Facility. This research was supported by the PsychENCODE Consortium (R01MH122509, H.W.), the IGVF Consortium (UM1HG012003, H.W.), the NICHD (5T32HD040127, J.L.), the NIH (MH132710, NS116859, E.A.), and BBRF Distinguished Investigator Award (E.A.).

## AUTHOR CONTRIBUTIONS

H.W., J.L. and E.S.A. designed the study and led the analysis. J.L. and A.T.L. conducted CROP-seq experiments. J.L., C.C., H.L., A.T.L., W.M. and Y.G. conducted and analyzed cilia imaging experiments. H.M. conducted the cNMF analysis. H.M., E.S.M., L.X. and M.J.Z. supported the establishment of the *in vivo* workflow. C.F., S.H.K., K.M., K.R.M. and S.C.H. provided and analyzed the human spatial transcriptomics data. J.L., H.M., A.T.L., E.S.A. and H.W. generated the figures. J.L., H.M., A.T.L., E.S.A. and H.W. co-wrote the manuscript, which was subsequently reviewed and edited by all authors.

## STAR METHODS

### EXPERIMENTAL MODEL DETAILS

All mouse experiments were conducted in accordance with the guidelines of the UNC-Chapel Hill Institutional Animal Care and Use Committee (IACUC), under the approved protocol no. 23-008.0. For CROP-seq experiments, postnatal day (P)13-15 heterozygous Cas9 transgenic mice^17^ (JAX, RRID:IMSR_JAX:026179) were used. Heterozygous mice were generated by crossing homozygous males to wild-type C57BL/6J females. After three days of mating, dams were separated from males and group-housed until 3-5 days before parturition. A total of 21 mice (10 males, 11 females) were used across 12 experimental batches, with each batch corresponding to one sequencing library. At the time of sacrifice, toes were collected from the pups for genotyping to confirm the presence of the Cas9 transgene. Genotyping was performed using PCR primers and protocols provided by the Jackson Laboratory. For cilia imaging experiments, P0 homozygous Cas9 transgenic pups from 4 litters were used. Homozygous mice were generated by crossing homozygous males to homozygous females.

## METHOD DETAILS

### CROP-seq target gene selection and gRNA design

We selected 12 genes mapped to SCZ risk variants that have empirically validated gene regulatory functions,^10^ and designed two 20-bp gRNAs per gene (**Table S1A**). For gRNA design, we used Benchling (RRID:SCR_013955). Annotated gene sequences from the mouse genome (mm10) were imported into Benchling, and gRNAs were selected from the 5’ region of the protein-coding sequence using an On-Target Score threshold of ≥65.

### CROP-seq library construction

We used the CROPseq-tRFP vector (Addgene, cat. no. #230938) for gRNA library construction. For each gRNA, a 74-bp single-stranded DNA oligonucleotide (ssDNA) was synthesized by Integrated DNA Technologies (IDT). Each oligonucleotide consisted of a 19-bp 5’ prefix (TGGAAAGGACGAAACACCG), a 20-bp gRNA sequence, and a 35-bp 3’ postfix (GTTTTAGAGCTAGAAATAGCAAGTTAAAATAAGGC), as previously described.^15^ The CROPseq-tRFP vector was digested with BsmB1 (NEB, cat. no. R0739S) at 55°C for 1.5 h followed by heat inactivation at 80°C for 20 min. To prevent re-ligation, the reaction was treated with recombinant shrimp alkaline phosphatase (rSAP; NEB, cat. no. M0371S) at 37°C for 1 h followed by heat inactivation at 65°C for 5 min. The digested product was resolved on a 0.8% agarose gel, and the 8,437-bp band was excised and purified using Zymoclean Gel DNA Recovery Kit (Zymo Research, cat. no. D4008). Next, Gibson assembly of the digested vector and gRNA oligonucleotides was performed using NEBuilder HiFi DNA Assembly Master Mix (NEB, cat. no. E2621S). All gRNA oligos were prepared at equal concentrations and pooled in equal volumes to generate a final pool concentration of 100 nM. Each reaction contained 11 fmol (57.35 ng) of digested vector and 200 fmol of pooled gRNA oligonucleotides. Assembly was carried out at 50 °C for 1 h, and the resulting library was purified using DNA Clean & Concentrator-5 kit (Zymo Research, cat. no. D4004). The assembled CROP-seq library was used to transform Endura Electrocompetent Cells (Lucigen, cat. no. 60242-2), followed by plating, culture, and maxiprep using ZymoPURE II Plasmid Maxiprep Kit (Zymo Research, cat. no. D4203). All culture media and plates contained carbenicillin (Thermo Scientific Chemicals, cat. no. AAJ6194906).

### CROP-seq library quality control

We used primers LibQC_i5 and LibQC_i7 (**Table S4A**) for PCR amplification of the gRNA region in the CROP-seq library, as previously described.^15^ LibQC_i7 primers included an 8-bp index sequence to label individual samples. To determine the optimal number of amplification cycles, we first performed quantitative PCR (qPCR) using NEBNext High-Fidelity 2X PCR Master Mix (NEB, cat. no. M0541) and SYBR Green I Nucleic Acid Gel Stain (Invitrogen, cat. no. S7563). Each reaction was performed with 10 ng of CROP-seq library DNA and primers at a final concentration of 0.5 µM, using an annealing temperature of 70 °C. The optimal cycle number was defined as the point at which the qPCR amplification curve reached one-quarter of the maximum fluorescence plateau. Subsequently, gRNA amplicons were generated using the determined number of cycles with four reactions. The four PCR reaction products were then pooled and purified using DNA Clean & Concentrator-5 kit (Zymo Research, cat. no. D4004). Amplicon size was verified using a TapeStation system (Agilent, cat. no. High Sensitivity D1000), with the expected product size of 281 bp. The amplicon was then sequenced on an Illumina MiSeq using a custom cycle configuration (Read 1 x Index = 50 x 8) with a 20% PhiX spike-in. Sequencing files were analyzed using custom scripts to assess the presence and relative abundance of each gRNA in the library.

### Lentiviral delivery of CROP-seq library to mouse brain

The CROP-seq library (CROPseq-tRFP-guideLib) was packaged into lentivirus by the UNC Lenti-shRNA Core Facility, yielding a titer of 2.25 x 10^9^ to 2.38 x 10^10^ transducing units (TU)/mL. On postnatal day 0, mouse pups were anesthetized on ice for 2 min prior to injection. Lentivirus mixed with Fast Green FCF dye (Sigma, cat. no. F7252-5G; final dye concentration, 0.5 mg/mL) was injected into the lateral ventricles of both hemispheres, with 1.5 µL injected per hemisphere. Injections were performed using a Hamilton syringe (Hamilton, cat. no. 7634-01) with a 32-gauge needle (Hamilton, cat. no. 7803-04; point style 2; 0.4-inch length) mounted on a custom stereotaxic instrument (Kopf, cat. nos. 176-61-SB-A, 1449-A, 1772-A). Following injection, the pups were warmed on a heating pad before being returned to the dam.

### Single-cell dissociation

Single-cell dissociation was performed on postnatal day 13-15. Following cervical dislocation, brains were extracted and coronally sectioned using an acrylic coronal brain matrix (Roboz Surgical, cat. no. AL1175) and razor blades (VWR, cat. no. 55411-050). The neocortex was dissected from the brain sections on ice in a dissection buffer consisting of Hibernate A Low Fluorescence (BrainBits, cat. no. HALF500), B-27 Plus Supplement (50x; Thermo Fisher, cat. no. A3582801), and Y-27632 ROCK inhibitor (final concentration of 10 µM; Selleck Chemicals, cat. no. S1049). The tissue was minced into small pieces, and cell dissociation was performed using the Worthington Papain Dissociation System (Worthington, cat. no. LK003153), following the manufacturer’s instructions. The EBSS buffer was equilibrated with 95% O2 5% CO2 prior to use.

The tissue was incubated in a buffer containing papain, DNase, and Y-27632 ROCK inhibitor at 37°C for 90 min in a 5% CO2 incubator with constant gentle agitation. Following enzymatic digestion, the tissue was gently triturated using a serological pipet until a homogenous cell suspension was achieved, with no visible large tissue fragments. Cells were filtered through a 70 μm cell strainer, and EBSS buffer, protease inhibitor, and DNase were added and gently mixed by inversion. Cells were pelleted at room temperature (RT) at 300g for 5 min and then resuspended in a buffer consisting of Hibernate A Low Fluorescence, B-27 Plus Supplement, 10% FBS (Avantor, cat. no. 97068-085), and Y-27632 ROCK inhibitor. Resuspension was performed by very gentle vortexing.

### Fluorescence activated cell sorting (FACS)

FACS was performed by the UNC Advanced Analytics Core. Live cells were first identified and selected using SYTOX Blue (Thermo Fisher, cat. no. 50-113-7613) and Pacific Blue Annexin V (Thermo Fisher, cat. no. NC9818309) dye, followed by sorting of the RFP+ population.

### Single-cell gene expression library and single-cell gRNA library preparation

Single-cell gene expression libraries were generated by the UNC Advanced Analytics Core using the 10x Genomics Chromium Next GEM Single Cell 3’ Kit v3.1 (Dual Index). gRNA reads were separately enriched from the cDNA using a two-step PCR protocol (PCR1 and PCR2) with custom-design primers (GuideEnrich_1, GuideEnrich_2, GuideEnrich_3, and GuideEnrich_4; **Table S4B**). Both PCR1 and PCR2 were performed using NEBNext Q5 Hot Start HiFi PCR Master Mix (NEB, cat. no. M0543), with final primer concentrations of 0.5 µM and an annealing temperature of 69°C. For PCR1, GuideEnrich_1 and GuideEnrich_2 primers were used. For each cDNA sample, five PCR reactions were prepared (using 1 ng of cDNA per reaction). One reaction was used in qPCR, where the optimal cycle number was defined as the point at which the qPCR amplification curve reached one-quarter of the maximum fluorescence plateau. The remaining four reactions were then used in standard PCR with the determined cycle number. These PCR reactions were pooled and purified using DNA Clean & Concentrator-5 kit (Zymo Research, cat. no. D4004). For PCR2, GuideEnrich_3 and GuideEnrich_4 primers were used, which included a 10-bp index sequence to label individual samples. As in PCR1, five reactions were prepared using the entire PCR1 product: one for qPCR and four for standard PCR. The pooled PCR2 product was size-selected and purified using sparQ PureMag Beads (Quantabio, cat. no. 95196) with a 0.5x → 0.7x protocol, selecting fragments in the 300-700 bp range.

### Sequencing

Single-cell gene expression libraries were sequenced on an Illumina NextSeq 2000 platform with P2 or P3 flow cells, using a Read 1 × i5 index × i7 index × Read 2 configuration of 28 × 10 × 10 × 90 cycles and 1% PhiX spike-in. gRNA libraries were sequenced on an Illumina NovaSeq 6000 platform with an SP flow cell, using a Read 1 × i5 index × i7 index × Read 2 configuration of 28 × 10 × 10 × 42 cycles and 20% PhiX spike-in.

### *Matr3*, *Scyl1*, and non-targeting control gRNA library construction and lentivirus packaging

The gRNA sequences targeting *Matr3* and *Scyl1* used for the CROP-seq experiments (**Table S4C**) were also used for the cilia imaging experiments. Additionally, three non-targeting control gRNA sequences^56^ were included (**Table S4C**). Library construction steps were identical to those described in the ‘CROP-seq library construction’ section up to the Gibson assembly step. However, gRNA oligonucleotides were not pooled for Gibson assembly; each gRNA oligonucleotide was cloned individually. The assembled gRNA plasmids were used to transform One Shot Stbl3 Chemically Competent E. coli (Thermo Fisher, cat. no. C737303), followed by plating. Individual colonies were picked, cultured for miniprep, and subjected to whole-plasmid sequencing. Only plasmids with a perfect match to the expected sequence were used for maxiprep. For maxiprep, plasmids containing gRNAs targeting the same gene (*Matr3*, *Scyl1*, or non-targeting control) were pooled. Endura Electrocompetent Cells (Lucigen, cat. no. 60242-2) were used for transformation, followed by plating, culture, and maxiprep using NucleoBond Xtra Maxi EF (Takara, cat. no. 740424.50). All culture media and plates contained carbenicillin (Thermo Scientific Chemicals, cat. no. AAJ6194906). Each gRNA library was packaged into lentivirus by the Boston Children’s Hospital Viral Core, yielding titers of 2.04 x 10^11^ GC/mL for *Matr3*, 4.39 x 10^11^ GC/mL for *Scyl1*, and 2.96 x 10^11^ GC/mL for non-targeting control.

### Mouse neocortex primary culture, lentivirus transduction, and immunohistochemical labeling of primary cilia

Cerebral cortices of P0 Cas9 homozygous transgenic mice were dissected and manually triturated in media consisting of Dulbecco’s Modified Eagle Medium (DMEM; Gibco, cat. no. 31053-028), 10% fetal bovine serum (FBS; VWR, cat. no. 97068085), and 1% penicillin/streptomycin (PS; Gibco, cat. no. 15140-122). 250,000 cells/well were then plated on 8-well glass bottom dish (Lab-TekII, ThermoFischer, cat. no. 154534) coated with poly-D-lysine/laminin (Sigma-Aldrich, cat. no. P6407-5MG/Sigma-Aldrich, cat. no. L2020-1MG) and cultured for 7 days in DMEM/10% FBS/1% PS media at 37°C, 5% CO2. Cortical cells were then infected with *Matr3*, *Scyl1*, and non-targeting control gRNA lentiviruses (prepared to 1 x 10^8^ GC/mL) with MOI of 10. Media were replaced with fresh media after 12-16 hours and subsequently every 3-4 days. 14 days after transduction, cells were washed once with phosphate-buffered saline (PBS) and fixed with 4% paraformaldehyde for 20 min at RT. Following fixation, cells were washed three times with PBS containing 0.1% Triton X-100 (PBST; 5 min per wash, RT) and then incubated in blocking buffer (0.2% gelatin, 300 mM NaCl, 0.3% Triton X-100 in 1x PBS) for 1 h at RT. Samples were subsequently incubated with primary antibody in blocking buffer for 48 h at 4°C. After three washes with PBST (5 min each, RT), cells were incubated with secondary antibody in blocking buffer for 1 h at RT. Samples were then washed three additional times with PBST (15 min each, RT). After the final wash, coverslips were mounted for imaging. The following primary antibody was used: Arl13b (1:1000, rabbit, Proteintech, cat. no. 17711-1-AP). The following secondary antibody was used: anti-rabbit 647 (1:1000, Invitrogen, cat. no. A21245). DAPI (1:1000, Invitrogen, cat. no. D21490) was used as a nuclear counterstain.

### Cilia imaging and quantification

Images of Arl13b-positive primary cilia in RFP-positive cells were acquired using a Zeiss 780 confocal microscope equipped with a 20x objective. We collected images from a total of 16 biological replicates, where each replicate represents at least one brain. Primary cilia length was measured by manually tracing the Arl13b signal using ImageJ Fiji (version 1.54P).^65^ Primary cilia shape was assessed by visual inspection and classified into three categories: smooth, irregular, and branched. For branched cilia, the length of the longer branch was measured from the ciliary base to the tip. A total of 449 control, 421 *Matr3*, and 459 *Scyl1* cells were analyzed.

## QUANTIFICATION AND STATISTICAL ANALYSIS

### Sequencing file preprocessing and data quality control

For each experimental batch, raw FASTQ files from single-cell gene expression and gRNA libraries were processed using 10x Genomics Cell Ranger^66^ (version 6.1.2) to output a gene x cell unique molecular identifier (UMI) count matrix and a gRNA x cell UMI count matrix. For reference transcriptome, mm10 (GENCODE vM23/Ensembl 98) built by 10x Genomics was used. The count matrices for each batch were loaded into a Seurat^67^ object (version 5.0.2) in R. Cells with ≥10% mitochondrial gene expression or cells with zero gRNA reads were removed. After that, filtered batches were integrated into a single Seurat object using log-normalization and RPCA method. The integrated object was used for downstream analyses.

### Cell type annotation

Cell type annotation was performed using the R package SingleR^68^ (version 2.2.0), with the Allen Brain Mouse Whole Cortex and Hippocampus 10x dataset^21^ used as a reference. To reduce computational cost, the reference dataset was downsampled to 50,000 cells. The raw gene x cell count matrix was used as input for cell type annotation. Cells that could not be confidently assigned to a cell type were removed. The 42 detected cell subtypes were regrouped into six major cell types: glutamatergic neurons, GABAergic neurons, astrocytes, microglia, oligodendrocytes, and vascular cells.

### Differential gene expression (DEG) analysis

The process for identifying DEGs is visually summarized in **Figure S2**. Raw UMI counts were used for modeling. For each perturbation, cells were categorized as control or perturbed. Control cells had zero reads for all gRNAs targeting the gene of interest, whereas perturbed cells had ≥3 reads for at least one of the two gRNAs targeting the gene of interest. For each perturbation, test genes were defined as genes detected in at least one cell in both the control and perturbed groups. For each test gene, we fitted a negative binomial generalized linear model using the following formula: gene ∼ gRNA + total gene reads + number of unique genes + total gRNA reads + number of unique gRNAs + batch + mitochondrial gene %, where the gRNA term is a categorical variable indicating the assignment of each cell as control or perturbed. For each perturbation, p-values for the gRNA term coefficients were adjusted for multiple testing using the Benjamini-Hochberg method. Genes with FDR<0.05 were selected as DEG candidates. To further validate DEG candidates, we performed permutation analysis. The gRNA labels (control/perturbed) were randomly shuffled across cells, and the same model was fitted. This process was repeated 1,000 times for each DEG candidate. A p-value was computed by comparing the observed gRNA coefficient to the distribution of coefficients from the permutations. Genes with permutation-based p-values<0.05 were defined as final DEGs.

### Cell type-specific expression analysis

The raw UMI count matrix was log-normalized using Seurat’s NormalizeData() function and used for cell type-specific expression analysis. The expression matrix was filtered to include only the gene set of interest (e.g., perturbation-specific DEGs or program genes) and partitioned by cell type. For each cell type, expression was first averaged across cells, followed by averaging across genes. The resulting cell type x gene set heatmap was scaled across cell types.

### Gene ontology (GO) analysis

Biological pathways enriched in a gene set of interest were identified using the R package gprofiler2^69^ (version 0.2.3) with the following command:

go <-gost(geneset, organism = “mmusculus”, ordered_query = FALSE, significant = TRUE, user_threshold = 0.05, correction_method = “fdr”, custom_bg = NULL, sources = c(“GO”, “KEGG”, “REAC”))

Resulting GO terms were filtered to include only those with a term size between 10 and 500. Redundant terms were reduced using REVIGO.^70^

### Expression of DEGs, cNMF programs, and regulons in postmortem brains from individuals with schizophrenia

We used a snRNA-seq dataset^28^ comprising 25 distinct cell types from the prefrontal cortex of individuals with SCZ for a permutation analysis. For each gene in a cell type-specific dataset, a differential expression (DE) score was defined as-log_10_P x logFC, where logFC represents the expression change in SCZ versus control, and P represents the significance of the change. Next, we identified the intersection between all genes in the cell type-specific dataset and all tested genes from the mouse CROP-seq data, converted to their human orthologs. From this intersection (universal background genes), N genes were randomly sampled 10,000 times, where N represents the number of mouse DEGs present in the universal background. A one-tailed p-value was computed by comparing the ∑(DE scores) for the DEGs to the distribution of ∑(DE scores) from the sampled genes. A p-value<0.05 indicated that the DEGs were significantly down-or upregulated in the given cell type in SCZ postmortem brain.

### Testing overlap between gene sets

We used Fisher’s exact tests to assess the significance of overlap between our gene sets (DEGs or regulon genes) and human risk gene sets. Mouse genes were first converted to human orthologs. The background gene set for each test was defined as the intersection of the background genes from the two datasets. Both datasets were restricted to genes within the corresponding background gene set.

### DEG program identification

We first constructed a DEG x perturbation matrix, where each entry represents the gRNA coefficient from the generalized linear model used in the differential gene expression analysis. From this matrix, we computed a DEG x DEG correlation matrix, where each entry represents the Spearman correlation coefficient between pairs of DEGs. The correlation matrix was converted to a distance matrix by subtracting each value from 1, and then subjected to hierarchical clustering. To determine the optimal number of clusters (k), we performed Silhouette score analysis and gene ontology analysis. When k = 5, the Silhouette score was the second highest, and the gene ontology results across the clusters were the most diverse with minimal redundancy. Therefore, we cut the hierarchical clustering tree at k = 5 to obtain five DEG programs.

### Regulation of DEG programs by perturbation

We constructed a DEG x perturbation matrix, where each entry represents the gRNA coefficient from the generalized linear model used in the differential gene expression analysis. This matrix was filtered to include only the genes within a given DEG program, followed by averaging across those genes to determine the average direction of regulation of the program by each perturbation.

### Consensus non-negative matrix factorization (cNMF)

To identify gene programs, we applied cNMF to the cell x gene count matrix, using the cNMF package^34^ (version 1.6.0; **Figure S4B**) We first performed CPM normalization and Harmony batch correction with the package functions.^71^ cNMF was then run across a range of factorization sizes (50-400 factors) using 2,000 highly variable genes, and results across different factorization sizes were systematically compared to select the optimal factorization size (**Figure S4D**).

### Gene assignment to cNMF factors

Genes were assigned to each factor based on their association strength with the corresponding factor. In previous studies using cNMF,^34,72^ a fixed number of top genes ranked by GEP (gene expression program) score are assigned to each factor regardless of their correlation strength with the factor. However, we observed that even genes within the top 300 GEP score may exhibit weak correlation with the corresponding factor (**Figure S4C**). Therefore, for each factor, we ranked genes by the absolute Spearman correlation between gene expression and factor scores (**Figure S4C**). Genes with |Spearman correlation|>0.1 were assigned to the factor; if fewer than 30 genes met this threshold, we additionally included top-ranked genes (allowing |Spearman correlation|≤0.1) until a minimum of 30 genes was reached. A maximum of 300 genes was assigned to each factor.

### Selection of the optimal factorization size

We repeated cNMF across factorization sizes ranging from 50 to 400. For each run, genes were assigned to factors as described above. Then across the factorization sizes, we compared (i) the size of the union of genes across factors and (ii) the size of the union of GO terms associated with these factors (**Figure S4D**). Based on these observations, we concluded that 300 factors sufficiently capture the full transcriptional architecture of the dataset, and used this factorization size for downstream analyses.

### Extraction of perturbation-associated general programs

To identify factors associated with perturbations, we performed Fisher’s exact tests between the genes assigned to each factor and the DEGs from each perturbation. Factors significantly associated with at least one perturbation (FDR<0.05) were selected, yielding a total of 118 factors (**Figure S4E**).

For biological interpretation, we further merged these factors into general programs through clustering (**Figure S4F**). We computed a Jaccard distance matrix between factor gene sets and performed hierarchical clustering using Ward’s linkage. Visualization using a t-SNE plot based on Jaccard distances confirmed eight clusters corresponding to eight general programs (**Figure S4F**). General program genes were defined as the union of genes across constituent subfactors. General program scores (**Figure 4C**) were defined as the mean score across constituent subfactors.

### Linear regression using cNMF program 0

A linear regression model (*P0 Score ∼ Perturbation category*) was used to test the effect of genetic perturbations on cNMF program 0 scores. Cells were grouped into four perturbation categories based on *Matr3* and *Scyl1* perturbation status (No pert, *Matr3*, *Scyl1*, Both). Using the fitted model, we then computed predicted P0 scores and corresponding confidence intervals for each perturbation category.

### Regulon identification using pySCENIC

We used the pySCENIC^35^ pipeline to identify regulons from the CROP-seq data. The mouse databases required to run the pipeline were downloaded from the Aerts Lab resource repository (https://resources.aertslab.org/). To improve computational efficiency, the input gene expression matrix was filtered to include only genes expressed in at least 1% of cells. The output regulons were filtered for enrichment in DEG program genes or cNMF program genes using a one-sided Fisher’s exact test, resulting in 167 regulons.

### Gene set relationship analysis

To summarize the relationships among the DEG programs, cNMF programs, and regulons, we constructed a gene set x gene set distance matrix, where each entry represents the distance (D) between a pair of gene sets. D was defined as 1/OR, where OR is the odds ratio from a one-sided Fisher’s exact test. This distance matrix was used for t-SNE dimensionality reduction. The resulting t-SNE coordinates were subjected to k-means clustering. The number of clusters (k) was selected such that each regulon was grouped with either a DEG or a cNMF program, and programs with similar biological functions were positioned in close proximity.

### Protein-protein interactions of regulon transcription factors and CROP-seq targets

The STRING^73^ database (version 12.0) was used to identify PPI interaction networks among regulon TFs and CROP-seq target genes. We used the full STRING network with an interaction score threshold of 0.7 (high-confidence). The resulting network edge list was downloaded and visualized with the igraph R package.^74^

### Identification of DEGs related to ciliary compartments and functions

We used the CilioGenics^45^ database, which provides scores representing the relevance of a given gene to cilia, and the SYSCILIA^46^ Gold Standard database, which annotates genes with their associated ciliary compartments. We identified the intersection between these two datasets and the DEGs converted to human orthologs, and then filtered the resulting gene set to include only those with a CilioGenics mean score ≥1.

### Comparison of CilioGenics scores of DEGs

Background genes were defined as all genes tested in the differential expression analysis. Gene symbols were converted to human orthologs for querying the CilioGenics^45^ database. Genes lacking a human ortholog or absent from the CilioGenics database were excluded. The “Neither” group comprised background genes that were neither *Matr3* nor *Scyl1* DEGs. Average CilioGenics scores were compared across the “Neither”, *Matr3* DEGs, and *Scyl1* DEGs using the Wilcoxon rank-sum test.

### Gene set enrichment analysis (GSEA)

GSEA was performed using the R package fgsea^75^ (version 1.32.4) with SCZ patient-derived gene sets from Kwon et al.^23^ The gene sets comprised all genes detected in 10x Genomics Visium-based spatially resolved transcriptomics (SRT) data from postmortem dorsolateral prefrontal cortex. Within each gene set, the genes were ranked by the t-statistic (t), which reflects the direction and significance of gene expression changes associated with high polygenic risk scores (PRS) or SCZ diagnosis. Positive t values (t>0) indicate increased gene expression with high PRS or SCZ, whereas negative t values (t<0) indicate decreased gene expression. Test gene sets consisted of *Matr3* and *Scyl1* DEGs converted to human orthologs. Enrichment of the test gene sets was assessed against the ranked SRT gene sets.

## Supplemental Figures

**Figure S1.**
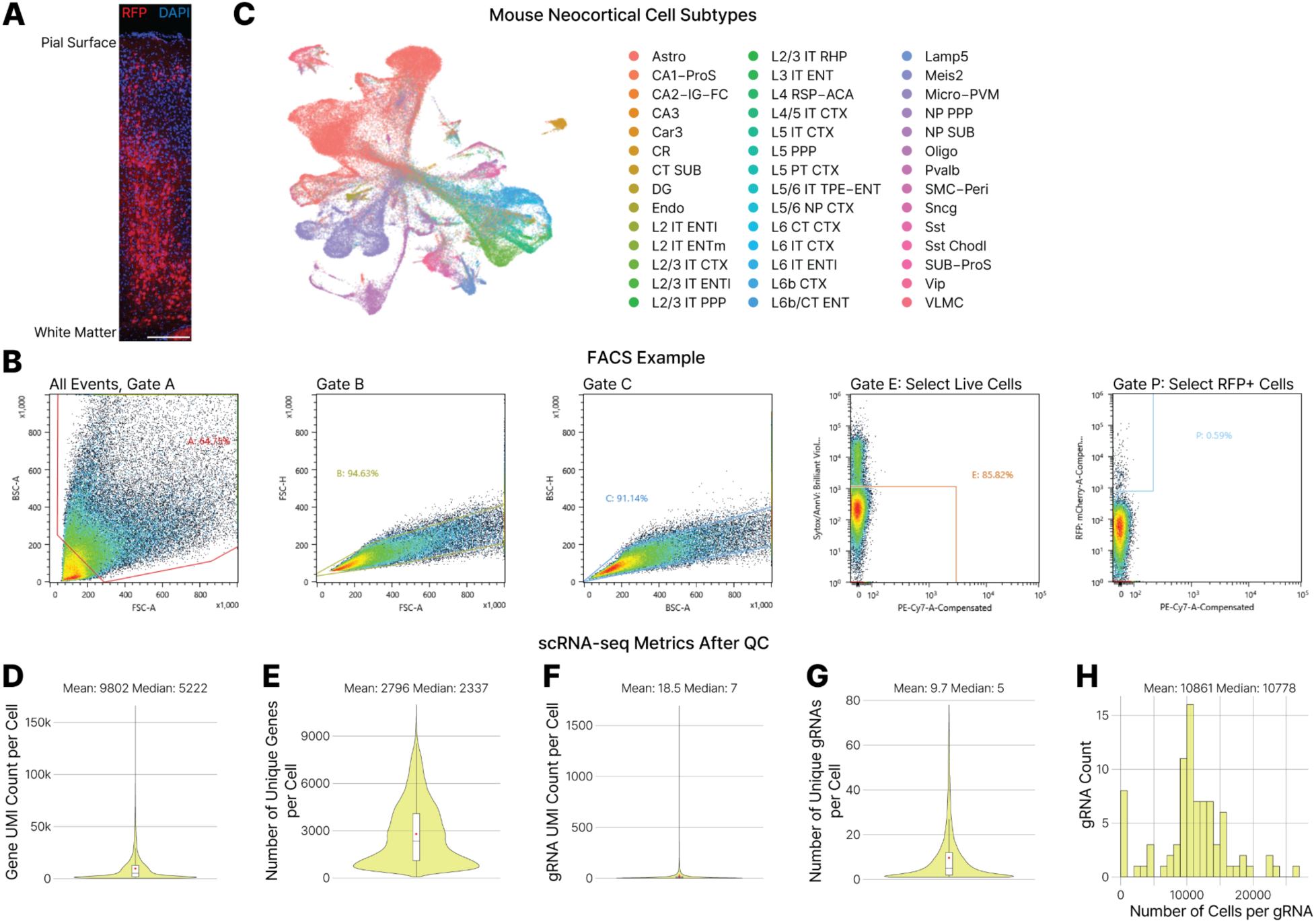
Collection and characterization of CROP-seq virus expressing cells. **A.** Expression of CROP-seq virus (RFP+) across the neocortical layers in the mouse brain. Scale bar, 200 μm. **B.** Example FACS gating steps to sort live RFP+ cells. **C.** Visualization of cell subtypes in the CROP-seq data. **D-E.** Distribution of per-cell gene UMI counts (**D**) and number of detected genes (**E**). **F-G.** Distribution of per-cell gRNA UMI counts (**F**) and number of detected gRNAs (**G**). **H.** Distribution of the number of cells per gRNA.

**Figure S2.**
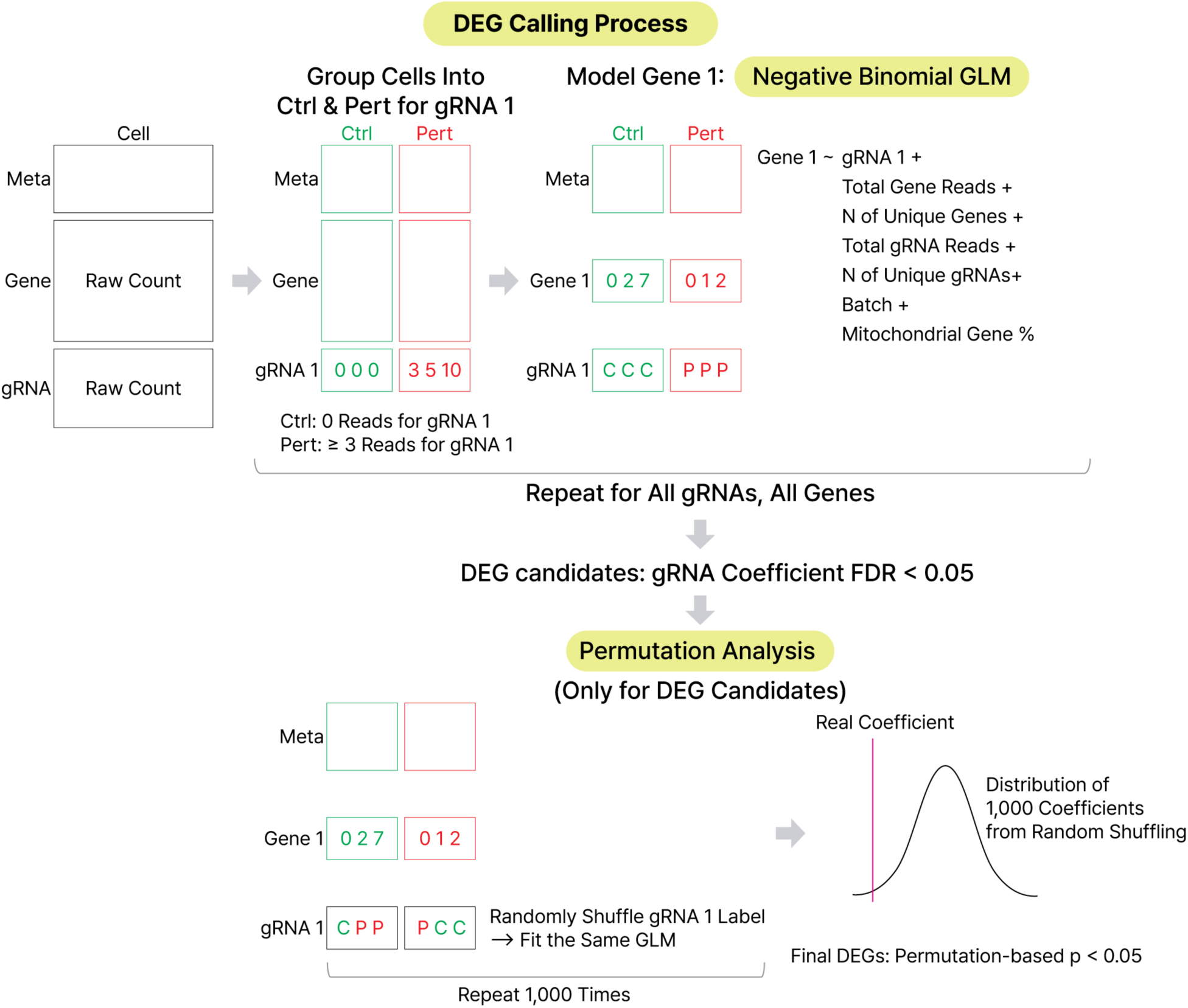
DEG calling workflow. For each perturbation, cells were grouped into control (Ctrl) and perturbed (Pert) populations based on gRNA detection. Differential expression was assessed using a negative binomial generalized linear model (GLM) that accounts for sequencing depth, gene detection rate, gRNA counts, batch, and mitochondrial gene content. Candidate DEGs (FDR<0.05) were further refined by permutation-based significance testing. See **Methods** for details.

**Figure S3.**
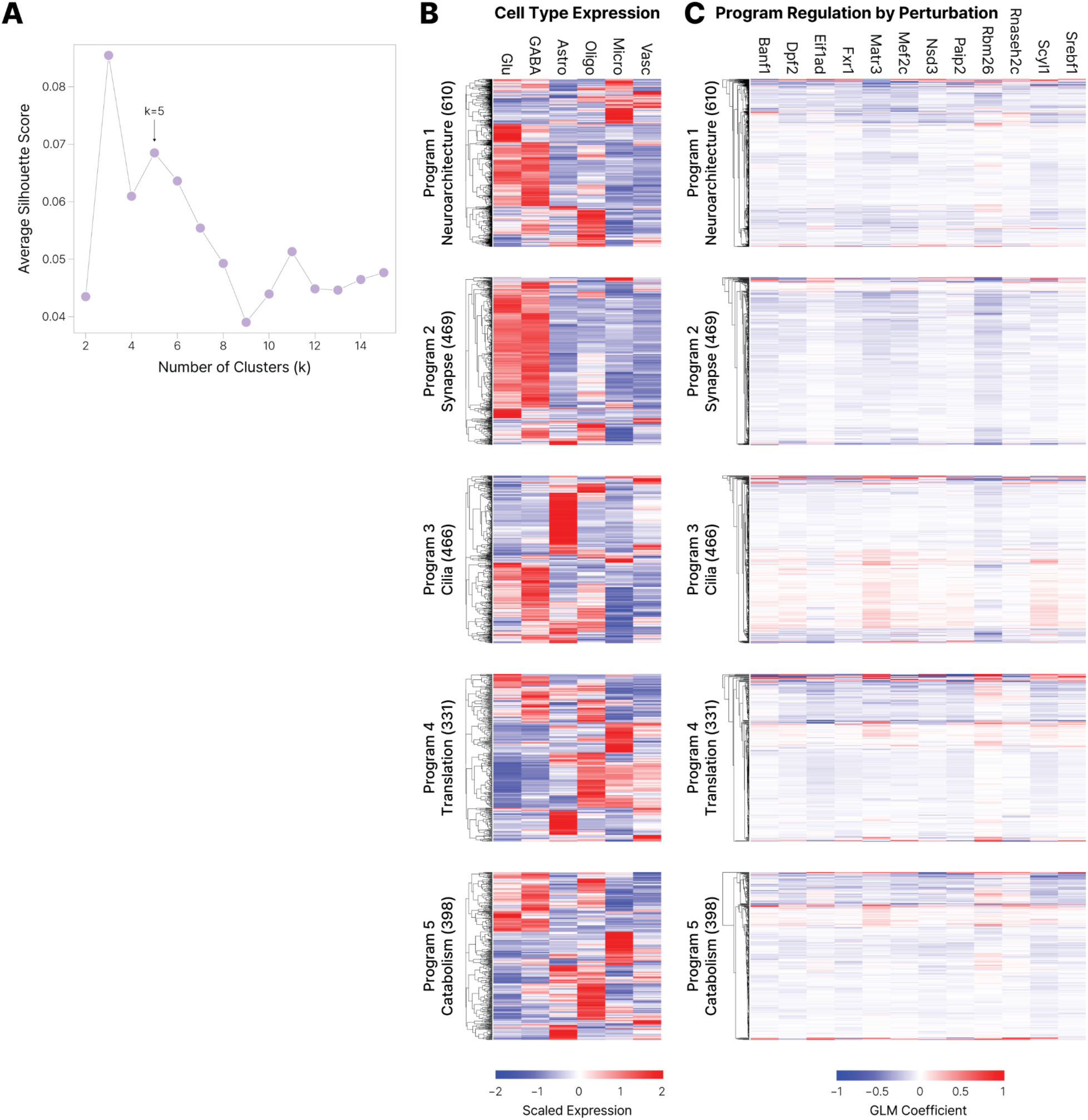
Characterization of DEG programs. **A.** Selection of the optimal number of clusters based on silhouette score, with k=5 chosen for downstream analysis. **B.** Cell type expression profiles of gene programs. Heatmaps show scaled expression of genes within each program across major cell types. **C.** Regulation of genes within each program across perturbations. Heatmaps show GLM coefficients for each gene across perturbations, indicating the direction and magnitude of differential expression.

**Figure S4.**
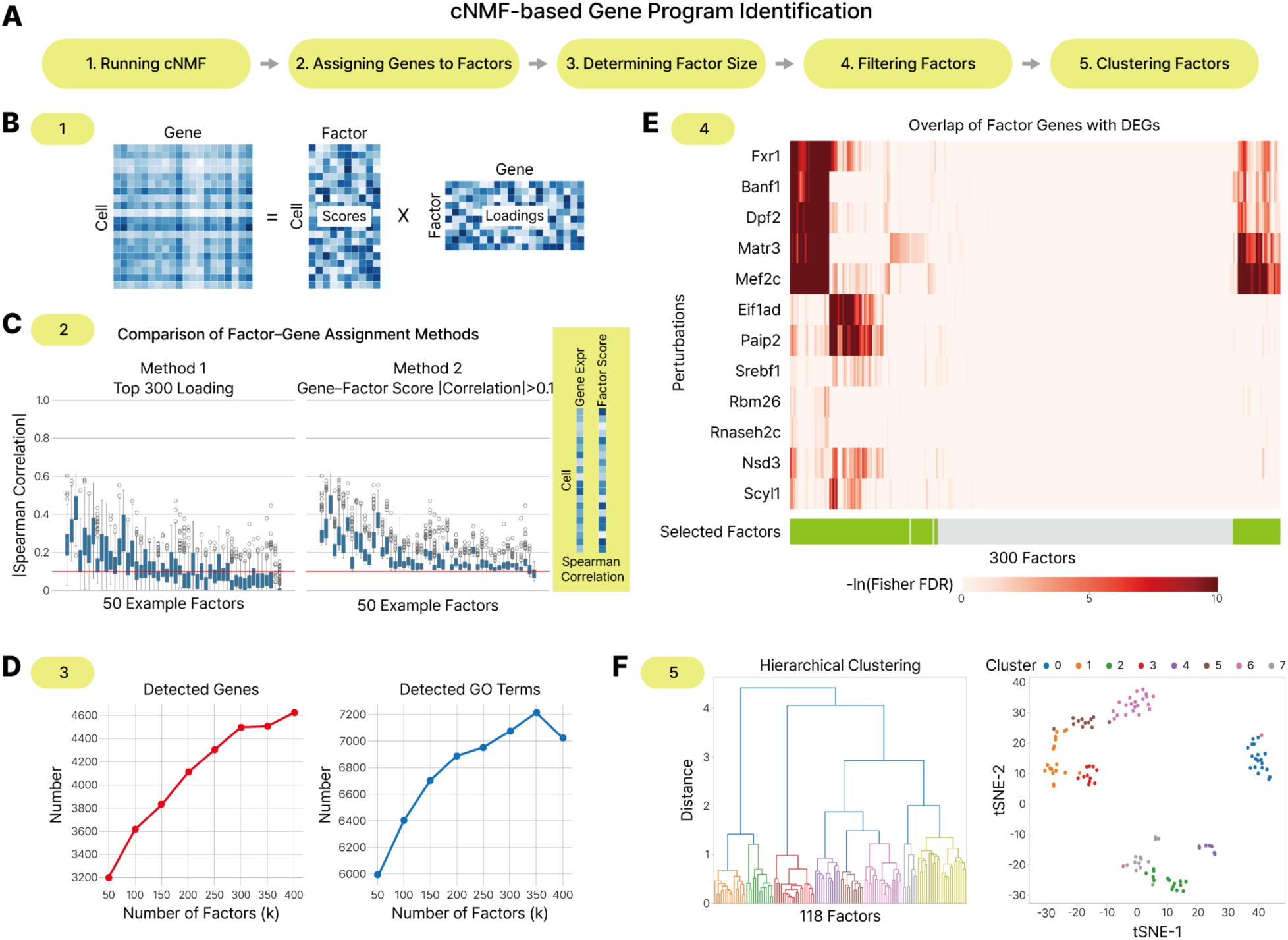
cNMF-based gene program identification workflow. **A.** Schematic overview of the cNMF program identification pipeline. **B.** Running cNMF. Decomposition of the single-cell expression matrix using cNMF into cell-factor score matrix and gene-factor loading matrix. **C.** Comparison of factor-gene assignment methods. Method 1 selects the top 300 genes based on cNMF loadings. Method 2 selects up to 300 genes with an absolute correlation>0.1 to cNMF factor scores. Box plots show the distribution of absolute Spearman correlation coefficients between gene expression and factor scores for the genes selected by each method. The red line indicates the absolute correlation threshold of 0.1. We used method 2 for downstream analyses because it more effectively selected genes with strong correlation to factor scores. **D.** Determining factor size (k). Total number of detected genes (left) and enriched GO (right) as a function of k. k=300 was selected based on plateauing trends in both metrics. **E.** Filtering factors. Overlap between 300 factors and perturbation-specific DEGs assessed by Fisher’s exact test. 118 factors were significantly associated with at least one perturbation (FDR<0.05). **F.** Clustering factors into gene programs. *Left*, hierarchical clustering of 118 factors into 8 programs. *Right*, t-SNE visualization of the 118 factors grouped into 8 programs.

**Figure S5.**
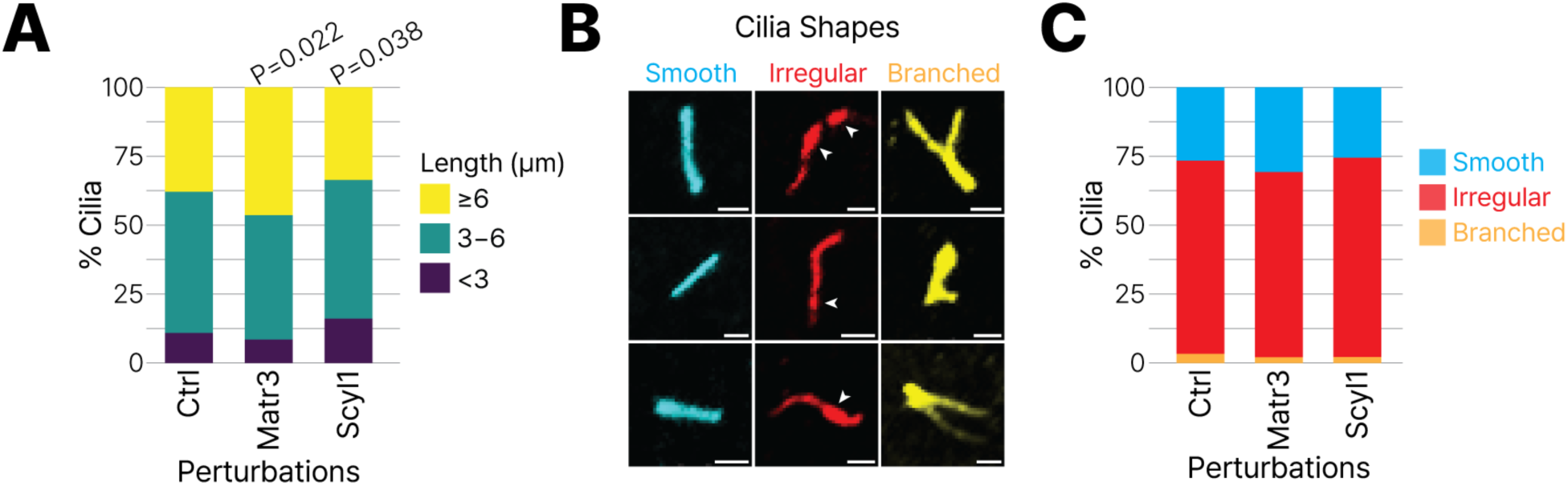
Effects of *Matr3* and *Scyl1* perturbations on primary cilia length and shape. **A.** Distribution of primary cilia lengths across perturbation groups, shown as the proportion of cilia in long (≥6 μm), medium (3-6 μm), and short (<3 μm) bins P-values indicate significant shifts in length distribution, as determined by a cumulative link mixed model. **B.** Representative examples of primary cilia shapes (smooth, irregular, branched) in mouse neocortical cells. Arrowheads indicate irregular, uneven, bead-like ciliary membrane domains. Scale bar, 2 μm. **C.** Distribution of cilia shape categories across perturbation groups, shown as the proportion of cilia classified as smooth, irregular, or branched.

## Supplemental Tables

**Table S1. Perturbations, gRNA sequences, and DEG analysis results**

**Table S2. DEG program, cNMF program, and regulon genes**

**Table S3. Ciliary DEGs**

**Table S4. Primer sequences and gRNA sequences for cilia imaging experiment**

## Key resources table

| REAGENT or RESOURCE | SOURCE | IDENTIFIER |
| --- | --- | --- |
| Antibodies |  |  |
| Rabbit anti-Arl13b | Proteintech | Cat# 17711-1-AP |
| Anti-rabbit 647 secondary antibody | Invitrogen | Cat# A21245 |
| Bacterial and virus strains |  |  |
| Endura Electrocompetent Cells | Lucigen | Cat# 60242-2 |
| One Shot Stbl3 Chemically Competent E. coli | Thermo Fisher | Cat# C737303 |
| CROPseq-tRFP-guideLib lentivirus | UNC Lenti-shRNA Core Facility | N/A |
| Matr3 gRNA lentivirus | Boston Children's Hospital Viral Core | N/A |
| Scyl1 gRNA lentivirus | Boston Children's Hospital Viral Core | N/A |
| Non-targeting control gRNA lentivirus | Boston Children's Hospital Viral Core | N/A |
| Chemicals, peptides, and recombinant proteins |  |  |
| BsmBI | NEB | Cat# R0739S |
| Recombinant shrimp alkaline phosphatase, rSAP | NEB | Cat# M0371S |
| NEBuilder HiFi DNA Assembly Master Mix | NEB | Cat# E2621S |
| Carbenicillin | Thermo Scientific Chemicals | Cat# AAJ6194906 |
| NEBNext High-Fidelity 2X PCR Master Mix | NEB | Cat# M0541 |
| SYBR Green I Nucleic Acid Gel Stain | Invitrogen | Cat# S7563 |
| Fast Green FCF dye | Sigma | Cat# F7252-5G |
| Hibernate A Low Fluorescence | BrainBits | Cat# HALF500 |
| B-27 Plus Supplement, 50x | Thermo Fisher | Cat# A3582801 |
| Y-27632 ROCK inhibitor | Selleck Chemicals | Cat# S1049 |
| Fetal bovine serum, FBS | Avantor/VWR | Cat# 97068085 |
| SYTOX Blue | Thermo Fisher | Cat# 50-113-7613 |
| Pacific Blue Annexin V | Thermo Fisher | Cat# NC9818309 |
| NEBNext Q5 Hot Start HiFi PCR Master Mix | NEB | Cat# M0543 |
| Dulbecco's Modified Eagle Medium, DMEM | Gibco | Cat# 31053-028 |
| Penicillin/Streptomycin | Gibco | Cat# 15140-122 |
| Poly-D-lysine | Sigma-Aldrich | Cat# P6407-5MG |
| Laminin | Sigma-Aldrich | Cat# L2020-1MG |
| DAPI | Invitrogen | Cat# D21490 |
| Critical commercial assays |  |  |
| Zymoclean Gel DNA Recovery Kit | Zymo Research | Cat# D4008 |
| DNA Clean & Concentrator-5 Kit | Zymo Research | Cat# D4004 |
| ZymoPURE II Plasmid Maxiprep Kit | Zymo Research | Cat# D4203 |
| TapeStation High Sensitivity D1000 | Agilent | Cat# High Sensitivity D1000 |
| Worthington Papain Dissociation System | Worthington | Cat# LK003153 |
| sparQ PureMag Beads | Quantabio | Cat# 95196 |
| NucleoBond Xtra Maxi EF | Takara | Cat# 740424.50 |
| Deposited data |  |  |
| Raw and processed CROP-seq data | This paper | GEO: GSE334303 |
| Codes for analyses | This paper | https://github.com/thewonlab/SCZ_CROPseq |
| Experimental models: Organisms/strains |  |  |
| Cas9 transgenic mice | The Jackson Laboratory | RRID:IMSR_JAX:026179 |
| Oligonucleotides |  |  |
| CROP-seq gRNA sequences in Table S1 |  |  |
| Primer sequences in Table S4 |  |  |
| Cilia imaging gRNA sequences in Table S4 |  |  |
| Recombinant DNA |  |  |
| CROPseq-tRFP vector | Addgene | Cat# 230938 |
| Software and algorithms |  |  |
| Benchling | Benchling | https://www.benchling.com/ |
| ImageJ Fiji (version 1.54P) | Schindelin et al.<sup>65</sup> | https://imagej.net/software/fiji/ |
| Cell Ranger (version 6.1.2) | Zheng et al.<sup>66</sup> | https://www.10xgenomics.com/support/software/cell-ranger/latest |
| Seurat (version 5.0.2) | Hao et al.<sup>67</sup> | https://satijalab.org/seurat/ |
| SingleR (version 2.2.0) | Aran et al.<sup>68</sup> | https://bioconductor.org/packages/release/bioc/html/SingleR.html |
| gprofiler2 (version 0.2.3) | Kolberg et al.<sup>69</sup> | https://biit.cs.ut.ee/gprofiler/page/r |
| REVIGO | Supek et al.<sup>70</sup> | https://revigo.irb.hr/ |
| cNMF (version 1.6.0) | Kotliar et al.<sup>34</sup> | https://github.com/dylkot/cNMF |
| pySCENIC | Aibar et al.<sup>35</sup> | https://pyscenic.readthedocs.io/en/latest/# |
| igraph | Antonov et al.<sup>74</sup> | https://r.igraph.org/articles/igraph.html |
| Other |  |  |
| Hamilton syringe | Hamilton | Cat# 7634-01 |
| 32-gauge Hamilton needle | Hamilton | Cat# 7803-04 |
| Custom stereotaxic instrument | Kopf | Cat# 176-61-SB-A, 1449-A, 1772-A |
| Acrylic coronal brain matrix | Roboz Surgical | Cat# AL1175 |
| Razor blades | VWR | Cat# 55411-050 |
| 8-well glass bottom dish, Lab-Tek II | Thermo Fisher | Cat# 154534 |

## REFERENCES

1. American Psychiatric Association (2013). Diagnostic and statistical manual of mental disorders (DSM-5 (R)) (American Psychiatric Association Publishing).

2. Sey, N.Y.A., Hu, B., Mah, W., Fauni, H., McAfee, J.C., Rajarajan, P., Brennand, K.J., Akbarian, S., and Won, H. (2020). A computational tool (H-MAGMA) for improved prediction of brain-disorder risk genes by incorporating brain chromatin interaction profiles. Nat. Neurosci. 23, 583–593.

3. Mah, W., and Won, H. (2020). The three-dimensional landscape of the genome in human brain tissue unveils regulatory mechanisms leading to schizophrenia risk. Schizophr. Res. 217, 17–25.

4. Sullivan, P.F., Kendler, K.S., and Neale, M.C. (2003). Schizophrenia as a complex trait: evidence from a meta-analysis of twin studies. Arch. Gen. Psychiatry 60, 1187–1192.

5. Trifu, S.C., Kohn, B., Vlasie, A., and Patrichi, B.-E. (2020). Genetics of schizophrenia (review). Exp. Ther. Med. 20, 3462–3468.

6. Trubetskoy, V., Pardiñas, A.F., Qi, T., Panagiotaropoulou, G., Awasthi, S., Bigdeli, T.B., Bryois, J., Chen, C.-Y., Dennison, C.A., Hall, L.S., et al. (2022). Mapping genomic loci implicates genes and synaptic biology in schizophrenia. Nature 604, 502–508.

7. Pardiñas, A.F., Holmans, P., Pocklington, A.J., Escott-Price, V., Ripke, S., Carrera, N., Legge, S.E., Bishop, S., Cameron, D., Hamshere, M.L., et al. (2018). Common schizophrenia alleles are enriched in mutation-intolerant genes and in regions under strong background selection. Nat. Genet. 50, 381–389.

8. Singh, T., Poterba, T., Curtis, D., Akil, H., Al Eissa, M., Barchas, J.D., Bass, N., Bigdeli, T.B., Breen, G., Bromet, E.J., et al. (2022). Rare coding variants in ten genes confer substantial risk for schizophrenia. Nature 604, 509–516.

9. Skene, N.G., Bryois, J., Bakken, T.E., Breen, G., Crowley, J.J., Gaspar, H.A., Giusti-Rodriguez, P., Hodge, R.D., Miller, J.A., Muñoz-Manchado, A.B., et al. (2018). Genetic identification of brain cell types underlying schizophrenia. Nat. Genet. 50, 825–833.

10. McAfee, J.C., Lee, S., Lee, J., Bell, J.L., Krupa, O., Davis, J., Insigne, K., Bond, M.L., Zhao, N., Boyle, A.P., et al. (2023). Systematic investigation of allelic regulatory activity of schizophrenia-associated common variants. Cell Genom. 3, 100404.

11. Jin, X., Simmons, S.K., Guo, A., Shetty, A.S., Ko, M., Nguyen, L., Jokhi, V., Robinson, E., Oyler, P., Curry, N., et al. (2020). In vivo Perturb-Seq reveals neuronal and glial abnormalities associated with autism risk genes. Science 370, eaaz6063.

12. Shi, T., Korshunova, M., Kim, S., DeTomaso, D., Zheng, X., Vishvanath, L., Nyasulu, T., Huynh, N., Sun, A., Thompson, P.C., et al. (2026). Genome-scale functional mapping of the mammalian whole brain with in vivo Perturb-seq. bioRxivorg. 10.64898/2026.03.16.711480.

13. Wu, B., Simmons, S.K., Kim, S., Li, J., Akram, M.A., Park, C.S., Zheng, X., Mendez, I., Patel, S., Chau, A., et al. (2026). Concordant transcriptional and morphological remodeling revealed by in vivo Perturb-CLEAR. bioRxivorg. 10.64898/2026.04.06.716787.

14. Santinha, A.J., Klingler, E., Kuhn, M., Farouni, R., Lagler, S., Kalamakis, G., Lischetti, U., Jabaudon, D., and Platt, R.J. (2023). Transcriptional linkage analysis with in vivo AAV-Perturb-seq. Nature 622, 367–375.

15. Datlinger, P., Rendeiro, A.F., Schmidl, C., Krausgruber, T., Traxler, P., Klughammer, J., Schuster, L.C., Kuchler, A., Alpar, D., and Bock, C. (2017). Pooled CRISPR screening with single-cell transcriptome readout. Nat. Methods 14, 297–301.

16. Lee, S., McAfee, J.C., Lee, J., Gomez, A., Ledford, A.T., Clarke, D., Min, H., Gerstein, M.B., Boyle, A.P., Sullivan, P.F., et al. (2025). Massively parallel reporter assay investigates shared genetic variants of eight psychiatric disorders. Cell 188, 1409–1424.e21.

17. Platt, R.J., Chen, S., Zhou, Y., Yim, M.J., Swiech, L., Kempton, H.R., Dahlman, J.E., Parnas, O., Eisenhaure, T.M., Jovanovic, M., et al. (2014). CRISPR-Cas9 knockin mice for genome editing and cancer modeling. Cell 159, 440–455.

18. Duncan, L.E., Li, T., Salem, M., Li, W., Mortazavi, L., Senturk, H., Shahverdizadeh, N., Vesuna, S., Shen, H., Yoon, J., et al. (2025). Mapping the cellular etiology of schizophrenia and complex brain phenotypes. Nat. Neurosci. 28, 248–258.

19. Song, L., Chen, W., Hou, J., Guo, M., and Yang, J. (2025). Spatially resolved mapping of cells associated with human complex traits. Nature 641, 932–941.

20. Smucny, J., Dienel, S.J., Lewis, D.A., and Carter, C.S. (2022). Mechanisms underlying dorsolateral prefrontal cortex contributions to cognitive dysfunction in schizophrenia. Neuropsychopharmacology 47, 292–308.

21. Yao, Z., van Velthoven, C.T.J., Nguyen, T.N., Goldy, J., Sedeno-Cortes, A.E., Baftizadeh, F., Bertagnolli, D., Casper, T., Chiang, M., Crichton, K., et al. (2021). A taxonomy of transcriptomic cell types across the isocortex and hippocampal formation. Cell 184, 3222–3241.e26.

22. Gandal, M.J., Zhang, P., Hadjimichael, E., Walker, R.L., Chen, C., Liu, S., Won, H., van Bakel, H., Varghese, M., Wang, Y., et al. (2018). Transcriptome-wide isoform-level dysregulation in ASD, schizophrenia, and bipolar disorder. Science 362, eaat8127.

23. Kwon, S.H., Guo, B., Fang, C., Tippani, M., Bach, S.V., Miller, R.A., Maguire, S.E., Iatrou, A., Pertea, G., Eagles, N.J., et al. (2026). Mapping spatially organized molecular and genetic signatures of schizophrenia across multiple scales in human prefrontal cortex. bioRxivorg. 10.64898/2026.02.16.706214.

24. Vergara, R.C., Jaramillo-Riveri, S., Luarte, A., Moënne-Loccoz, C., Fuentes, R., Couve, A., and Maldonado, P.E. (2019). The Energy Homeostasis Principle: Neuronal energy regulation drives local network dynamics generating behavior. Front. Comput. Neurosci. 13, 49.

25. Ni, P., Ma, Y., and Chung, S. (2024). Mitochondrial dysfunction in psychiatric disorders. Schizophr. Res. 273, 62–77.

26. Helaly, A.M.N., and Ghorab, D.S.E.D. (2023). Schizophrenia as metabolic disease. What are the causes? Metab. Brain Dis. 38, 795–804.

27. Fizíková, I., Dragašek, J., and Račay, P. (2023). Mitochondrial dysfunction, altered mitochondrial oxygen, and energy metabolism associated with the pathogenesis of schizophrenia. Int. J. Mol. Sci. 24. 10.3390/ijms24097991.

28. Ruzicka, W.B., Mohammadi, S., Fullard, J.F., Davila-Velderrain, J., Subburaju, S., Tso, D.R., Hourihan, M., Jiang, S., Lee, H.-C., Bendl, J., et al. (2024). Single-cell multi-cohort dissection of the schizophrenia transcriptome. Science 384, eadg5136.

29. Ling, E., Nemesh, J., Goldman, M., Kamitaki, N., Reed, N., Handsaker, R.E., Genovese, G., Vogelgsang, J.S., Gerges, S., Kashin, S., et al. (2024). A concerted neuron-astrocyte program declines in ageing and schizophrenia. Nature 627, 604–611.

30. Kaplanis, J., Samocha, K.E., Wiel, L., Zhang, Z., Arvai, K.J., Eberhardt, R.Y., Gallone, G., Lelieveld, S.H., Martin, H.C., McRae, J.F., et al. (2020). Evidence for 28 genetic disorders discovered by combining healthcare and research data. Nature 586, 757–762.

31. Replogle, J.M., Saunders, R.A., Pogson, A.N., Hussmann, J.A., Lenail, A., Guna, A., Mascibroda, L., Wagner, E.J., Adelman, K., Lithwick-Yanai, G., et al. (2022). Mapping information-rich genotype-phenotype landscapes with genome-scale Perturb-seq. Cell 185, 2559–2575.e28.

32. Wang, Y., Armendariz, D.A., Wang, L., Zhao, H., Xie, S., and Hon, G.C. (2025). Enhancer regulatory networks globally connect non-coding breast cancer loci to cancer genes. Genome Biol. 26, 10.

33. Li, C., Fleck, J.S., Martins-Costa, C., Burkard, T.R., Themann, J., Stuempflen, M., Peer, A.M., Vertesy, Á., Littleboy, J.B., Esk, C., et al. (2023). Single-cell brain organoid screening identifies developmental defects in autism. Nature 621, 373–380.

34. Kotliar, D., Veres, A., Nagy, M.A., Tabrizi, S., Hodis, E., Melton, D.A., and Sabeti, P.C. (2019). Identifying gene expression programs of cell-type identity and cellular activity with single-cell RNA-Seq. Elife 8. 10.7554/eLife.43803.

35. Aibar, S., González-Blas, C.B., Moerman, T., Huynh-Thu, V.A., Imrichova, H., Hulselmans, G., Rambow, F., Marine, J.-C., Geurts, P., Aerts, J., et al. (2017). SCENIC: single-cell regulatory network inference and clustering. Nat. Methods 14, 1083–1086.

36. Fu, J.M., Satterstrom, F.K., Peng, M., Brand, H., Collins, R.L., Dong, S., Wamsley, B., Klei, L., Wang, L., Hao, S.P., et al. (2022). Rare coding variation provides insight into the genetic architecture and phenotypic context of autism. Nat. Genet. 54, 1320–1331.

37. Choksi, S.P., Lauter, G., Swoboda, P., and Roy, S. (2014). Switching on cilia: transcriptional networks regulating ciliogenesis. Development 141, 1427–1441.

38. Coyle, M.C., Tajima, A.M., Leon, F., Choksi, S.P., Yang, A., Espinoza, S., Hughes, T.R., Reiter, J.F., Booth, D.S., and King, N. (2023). An RFX transcription factor regulates ciliogenesis in the closest living relatives of animals. Curr. Biol. 33, 3747–3758.e9.

39. Wu, J.Y., Cho, S.-J., Descant, K., Li, P.H., Shapson-Coe, A., Januszewski, M., Berger, D.R., Meyer, C., Casingal, C., Huda, A., et al. (2024). Mapping of neuronal and glial primary cilia contactome and connectome in the human cerebral cortex. Neuron 112, 41–55.e3.

40. Loukil, A., Ebright, E., Hamdi, K., Menzel, E., Uezu, A., Gao, Y., Soderling, S.H., and Goetz, S.C. (2025). Identification of new ciliary signaling pathways in the brain and insights into neurological disorders. J. Neurosci. 45, e0800242025.

41. Hilgendorf, K.I., Johnson, C.T., and Jackson, P.K. (2016). The primary cilium as a cellular receiver: organizing ciliary GPCR signaling. Curr. Opin. Cell Biol. 39, 84–92.

42. Kostyanovskaya, E., Lasser, M.C., Wang, B., Schmidt, J., Bader, E., Buteo, C., Arbelaez, J., Sindledecker, A.R., McCluskey, K.E., Castillo, O., et al. (2025). Convergence of autism proteins at the cilium. bioRxivorg. 10.1101/2024.12.05.626924.

43. Alhassen, W., Chen, S., Vawter, M., Robbins, B.K., Nguyen, H., Myint, T.N., Saito, Y., Schulmann, A., Nauli, S.M., Civelli, O., et al. (2021). Patterns of cilia gene dysregulations in major psychiatric disorders. Prog. Neuropsychopharmacol. Biol. Psychiatry 109, 110255.

44. Muñoz-Estrada, J., Lora-Castellanos, A., Meza, I., Alarcón Elizalde, S., and Benítez-King, G. (2018). Primary cilia formation is diminished in schizophrenia and bipolar disorder: A possible marker for these psychiatric diseases. Schizophr. Res. 195, 412–420.

45. Pir, M.S., Begar, E., Yenisert, F., Demirci, H.C., Korkmaz, M.E., Karaman, A., Tsiropoulou, S., Firat-Karalar, E.N., Blacque, O.E., Oner, S.S., et al. (2024). CilioGenics: an integrated method and database for predicting novel ciliary genes. Nucleic Acids Res. 52, 8127–8145.

46. Vasquez, S.S.V., van Dam, J., and Wheway, G. (2021). An updated SYSCILIA gold standard (SCGSv2) of known ciliary genes, revealing the vast progress that has been made in the cilia research field. Mol. Biol. Cell 32, br13.

47. Santos, J.R., and Park, J. (2024). MATR3’s role beyond the nuclear matrix: From gene regulation to its implications in amyotrophic lateral sclerosis and other diseases. Cells 13, 980.

48. Rajarajan, P., Borrman, T., Liao, W., Schrode, N., Flaherty, E., Casiño, C., Powell, S., Yashaswini, C., LaMarca, E.A., Kassim, B., et al. (2018). Neuron-specific signatures in the chromosomal connectome associated with schizophrenia risk. Science 362, eaat4311.

49. Pelletier, S. (2016). SCYL pseudokinases in neuronal function and survival. Neural Regen. Res. 11, 42–44.

50. Kaeser-Pebernard, S., Vionnet, C., Mari, M., Sankar, D.S., Hu, Z., Roubaty, C., Martínez-Martínez, E., Zhao, H., Spuch-Calvar, M., Petri-Fink, A., et al. (2022). mTORC1 controls Golgi architecture and vesicle secretion by phosphorylation of SCYL1. Nat. Commun. 13, 4685.

51. Stevenson, N.L. (2023). The factory, the antenna and the scaffold: the three-way interplay between the Golgi, cilium and extracellular matrix underlying tissue function. Biol. Open 12. 10.1242/bio.059719.

52. Riether, C., Radpour, R., Kallen, N.M., Bürgin, D.T., Bachmann, C., Schürch, C.M., Lüthi, U., Arambasic, M., Hoppe, S., Albers, C.E., et al. (2021). Metoclopramide treatment blocks CD93-signaling-mediated self-renewal of chronic myeloid leukemia stem cells. Cell Rep. 34, 108663.

53. Schmidt, W.M., Rutledge, S.L., Schüle, R., Mayerhofer, B., Züchner, S., Boltshauser, E., and Bittner, R.E. (2015). Disruptive SCYL1 mutations underlie a syndrome characterized by recurrent episodes of liver failure, peripheral neuropathy, cerebellar atrophy, and ataxia. Am. J. Hum. Genet. 97, 855–861.

54. Lenz, D., McClean, P., Kansu, A., Bonnen, P.E., Ranucci, G., Thiel, C., Straub, B.K., Harting, I., Alhaddad, B., Dimitrov, B., et al. (2018). SCYL1 variants cause a syndrome with low γ-glutamyl-transferase cholestasis, acute liver failure, and neurodegeneration (CALFAN). Genet. Med. 20, 1255–1265.

55. Kuliyev, E., Gingras, S., Guy, C.S., Howell, S., Vogel, P., and Pelletier, S. (2018). Overlapping role of SCYL1 and SCYL3 in maintaining motor neuron viability. J. Neurosci. 38, 2615–2630.

56. Jost, M., Santos, D.A., Saunders, R.A., Horlbeck, M.A., Hawkins, J.S., Scaria, S.M., Norman, T.M., Hussmann, J.A., Liem, C.R., Gross, C.A., et al. (2020). Titrating gene expression using libraries of systematically attenuated CRISPR guide RNAs. Nat. Biotechnol. 38, 355–364.

57. Liu, S., Trupiano, M.X., Simon, J., Guo, J., and Anton, E.S. (2021). The essential role of primary cilia in cerebral cortical development and disorders. Curr. Top. Dev. Biol. 142, 99–146.

58. Macarelli, V., Leventea, E., and Merkle, F.T. (2023). Regulation of the length of neuronal primary cilia and its potential effects on signalling. Trends Cell Biol. 33, 979–990.

59. Volos, P., Fujise, K., and Rafiq, N.M. (2025). Roles for primary cilia in synapses and neurological disorders. Trends Cell Biol. 35, 6–10.

60. Kim, M., Haney, J.R., Zhang, P., Hernandez, L.M., Wang, L.-K., Perez-Cano, L., Loohuis, L.M.O., de la Torre-Ubieta, L., and Gandal, M.J. (2021). Brain gene co-expression networks link complement signaling with convergent synaptic pathology in schizophrenia. Nat. Neurosci. 24, 799–809.

61. Guo, J., Otis, J.M., Higginbotham, H., Monckton, C., Cheng, J., Asokan, A., Mykytyn, K., Caspary, T., Stuber, G.D., and Anton, E.S. (2017). Primary cilia signaling shapes the development of interneuronal connectivity. Dev. Cell 42, 286–300.e4.

62. Kumamoto, N., Gu, Y., Wang, J., Janoschka, S., Takemaru, K.-I., Levine, J., and Ge, S. (2012). A role for primary cilia in glutamatergic synaptic integration of adult-born neurons. Nat. Neurosci. 15, 399–405, S1.

63. Tereshko, L., Gao, Y., Cary, B.A., Turrigiano, G.G., and Sengupta, P. (2021). Ciliary neuropeptidergic signaling dynamically regulates excitatory synapses in postnatal neocortical pyramidal neurons. Elife 10. 10.7554/eLife.65427.

64. Sheu, S.-H., Upadhyayula, S., Dupuy, V., Pang, S., Deng, F., Wan, J., Walpita, D., Pasolli, H.A., Houser, J., Sanchez-Martinez, S., et al. (2022). A serotonergic axon-cilium synapse drives nuclear signaling to alter chromatin accessibility. Cell 185, 3390–3407.e18.

65. Schindelin, J., Arganda-Carreras, I., Frise, E., Kaynig, V., Longair, M., Pietzsch, T., Preibisch, S., Rueden, C., Saalfeld, S., Schmid, B., et al. (2012). Fiji: an open-source platform for biological-image analysis. Nat. Methods 9, 676–682.

66. Zheng, G.X.Y., Terry, J.M., Belgrader, P., Ryvkin, P., Bent, Z.W., Wilson, R., Ziraldo, S.B., Wheeler, T.D., McDermott, G.P., Zhu, J., et al. (2017). Massively parallel digital transcriptional profiling of single cells. Nat. Commun. 8, 14049.

67. Hao, Y., Stuart, T., Kowalski, M.H., Choudhary, S., Hoffman, P., Hartman, A., Srivastava, A., Molla, G., Madad, S., Fernandez-Granda, C., et al. (2024). Dictionary learning for integrative, multimodal and scalable single-cell analysis. Nat. Biotechnol. 42, 293–304.

68. Aran, D., Looney, A.P., Liu, L., Wu, E., Fong, V., Hsu, A., Chak, S., Naikawadi, R.P., Wolters, P.J., Abate, A.R., et al. (2019). Reference-based analysis of lung single-cell sequencing reveals a transitional profibrotic macrophage. Nat. Immunol. 20, 163–172.

69. Kolberg, L., Raudvere, U., Kuzmin, I., Vilo, J., and Peterson, H. (2020). gprofiler2 -- an R package for gene list functional enrichment analysis and namespace conversion toolset g:Profiler. F1000Res. 9, 709.

70. Supek, F., Bošnjak, M., Škunca, N., and Šmuc, T. (2011). REVIGO summarizes and visualizes long lists of gene ontology terms. PLoS One 6, e21800.

71. Kotliar, D., Curtis, M., Agnew, R., Weinand, K., Nathan, A., Baglaenko, Y., Slowikowski, K., Zhao, Y., Sabeti, P.C., Rao, D.A., et al. (2025). Reproducible single-cell annotation of programs underlying T cell subsets, activation states and functions. Nat. Methods 22, 1964–1980.

72. Schnitzler, G.R., Kang, H., Fang, S., Angom, R.S., Lee-Kim, V.S., Ma, X.R., Zhou, R., Zeng, T., Guo, K., Taylor, M.S., et al. (2024). Convergence of coronary artery disease genes onto endothelial cell programs. Nature 626, 799–807.

73. Szklarczyk, D., Kirsch, R., Koutrouli, M., Nastou, K., Mehryary, F., Hachilif, R., Gable, A.L., Fang, T., Doncheva, N.T., Pyysalo, S., et al. (2023). The STRING database in 2023: protein-protein association networks and functional enrichment analyses for any sequenced genome of interest. Nucleic Acids Res. 51, D638–D646.

74. Antonov, M., Csárdi, G., Horvát, S., Müller, K., Nepusz, T., Noom, D., Salmon, M., Traag, V., Welles, B.F., and Zanini, F. (2023). Igraph enables fast and robust network analysis across programming languages. arXiv [cs.SI]. 10.48550/arXiv.2311.10260.

75. Korotkevich, G., Sukhov, V., Budin, N., Shpak, B., Artyomov, M.N., and Sergushichev, A. (2016). Fast gene set enrichment analysis. bioRxiv. 10.1101/060012.

